# A Phosphoproteomics Data Resource for Systems-level Modeling of Kinase Signaling Networks

**DOI:** 10.1101/2023.08.03.551714

**Authors:** Song Feng, James A. Sanford, Thomas Weber, Chelsea M. Hutchinson-Bunch, Panshak P. Dakup, Vanessa L. Paurus, Kwame Attah, Herbert M. Sauro, Wei-Jun Qian, H. Steven Wiley

## Abstract

Building mechanistic models of kinase-driven signaling pathways requires quantitative measurements of protein phosphorylation across physiologically relevant conditions, but this is rarely done because of the insensitivity of traditional technologies. By using a multiplexed deep phosphoproteome profiling workflow, we were able to generate a deep phosphoproteomics dataset of the EGFR-MAPK pathway in non-transformed MCF10A cells across physiological ligand concentrations with a time resolution of <12 min and in the presence and absence of multiple kinase inhibitors. An improved phosphosite mapping technique allowed us to reliably identify >46,000 phosphorylation sites on >6600 proteins, of which >4500 sites from 2110 proteins displayed a >2-fold increase in phosphorylation in response to EGF. This data was then placed into a cellular context by linking it to 15 previously published protein databases. We found that our results were consistent with much, but not all previously reported data regarding the activation and negative feedback phosphorylation of core EGFR-ERK pathway proteins. We also found that EGFR signaling is biphasic with substrates downstream of RAS/MAPK activation showing a maximum response at <3ng/ml EGF while direct substrates, such as HGS and STAT5B, showing no saturation. We found that RAS activation is mediated by at least 3 parallel pathways, two of which depend on PTPN11. There appears to be an approximately 4-minute delay in pathway activation at the step between RAS and RAF, but subsequent pathway phosphorylation was extremely rapid. Approximately 80 proteins showed a >2-fold increase in phosphorylation across all experiments and these proteins had a significantly higher median number of phosphorylation sites (~18) relative to total cellular phosphoproteins (~4). Over 60% of EGF-stimulated phosphoproteins were downstream of MAPK and included mediators of cellular processes such as gene transcription, transport, signal transduction and cytoskeletal arrangement. Their phosphorylation was either linear with respect to MAPK activation or biphasic, corresponding to the biphasic signaling seen at the level of the EGFR. This deep, integrated phosphoproteomics data resource should be useful in building mechanistic models of EGFR and MAPK signaling and for understanding how downstream responses are regulated.

## Introduction

Signal transduction networks are crucial regulators of cell behavior, and their dysregulation is causal to many diseases, such as cancer (Sanchez-Vega et al. 2018). Because of their inherent complexity, much effort has focused on building predictive computational models of these networks that can predict not only how genetic alterations cause dysregulation, but which modifications or drug treatments can restore their normal function (Frohlich et al. 2023). Creating such models has proven to be very challenging because of both their complexity and the lack of suitable technologies to parameterize them. Theoretical studies have suggested that the number of measurements needed to constrain relatively simple models can be quite high (Apgar et al. 2010). Thus, most current models are either purposely simple or cannot be experimentally validated, either of which limits their usefulness.

Phosphoproteomics is becoming an increasingly powerful technology for generating molecular-level data on changes in protein activity and state in response to signaling and metabolic events (Pan et al. 2009; Humphrey et al. 2015; Yi et al. 2018). Because of both the precision and high-throughput nature of mass spectrometry, it can generate the quality and amount of data needed to build and constrain more realistic models of signaling networks. Biochemical studies have identified specific phosphorylation events that can either activate or inhibit signal transduction, thus linking an observed change in protein phosphorylation to alterations in signal flow within a pathway (Zheng et al. 2013; Lavoie and Therrien 2015). To make such an association, however, requires measuring the phosphorylation state of proteins under conditions that span the response profile of the cells. This, unfortunately, has rarely been done.

One of the most popular systems used for both modeling and phosphoproteomics studies is the EGFR-MAPK pathway (Reddy et al. 2016; Rudolph et al. 2016; Yi et al. 2018). The EGFR is a widely expressed receptor type possessing intrinsic tyrosine kinase activity. It is involved in a wide range of physiological processes and plays a causal role in multiple types of cancer (Hohensee et al. 2013; Orton et al. 2009; Sever and Brugge 2015). Because its kinase activity is central to its biological activity, numerous studies have tried to define the pattern of protein phosphorylation induced by EGFR activation and identify which phosphorylation events are responsible for its multiple physiological effects (Chen et al. 1989; Batzer et al. 1994; Keilhack et al. 1998; Gill et al. 2017). Previous phosphoproteomics datasets, however, were mostly generated from physiologically questionable systems, such as HeLa or HEK293 cells (Sharma et al. 2014; Hsu et al. 2011) and using non-physiological conditions, such as supraphysiological concentrations of EGF (>100ng/ml) (Blagoev et al. 2004) or using cells expressing extraordinarily high levels of receptors (Koksal et al. 2018; Foerster et al. 2013). These experimental design choices were likely made because of the relative insensitivities of the proteomics technologies that were available at the time. Unfortunately, cell lines such as HeLa have been selected for many decades for rapid in vitro growth and many of their key physiological functions have lost their dependence on signaling pathways, such as those regulated by EGF (Buvinic et al. 2007). Maximal activation of the MAPK pathway also occurs at <3ng/ml EGF (Pinilla-Macua et al. 2017), a level that produced barely detectible phosphorylation changes in previous studies (Reddy et al. 2016; Bisson et al. 2011). However, high concentrations of EGFR can activate many additional signaling pathways (Krall et al. 2011), thus complicating attempts to identify MAPK pathway-specific phosphorylation events. The lack of clear relationship between treatment regimens and downstream biological responses of previous studies has limited the usefulness of much of their data.

Recent advances in mass spectrometry-based proteomics have significantly increased both the sensitivity and coverage of phosphoproteome analyses, enabling the generation of robust and detailed quantitative datasets under more biologically relevant conditions (Humphrey et al. 2015). Particularly powerful are multiplexing techniques based on isobaric tags that allow multiple samples across time and dose to be simultaneously analyzed in a single multiplexed assay, while reducing sample-to-sample variation and noise (Rauniyar and Yates 2014; Mertins et al. 2018). However, to be most useful, data should be generated from widely used model systems that display well-characterized responses. In addition, the treatment time and dose should span relevant biological conditions. Towards this end, we sought to generate a comprehensive time-series, dose-response, and inhibitor-response phosphoproteomics dataset using the well-established MCF10A mammary epithelial cell line (Tait et al. 1990; Debnath et al. 2003). These non-cancer cells are normally dependent on activation of the EGFR system for their growth and have been used for measuring and modeling signaling pathways at both the population and single cell level (Gillies et al. 2017).

We first identified the most appropriate treatment conditions for our analyses using ELISA assays to quantify both RAS and MAPK activation. Using conditions that spanned the entire dose-time regime for RAS and MAPK activation (100pM-10nM, 1-12min), we generated a deep phosphoproteomics dataset that describes in new detail how cellular patterns of protein phosphorylation change in response to EGF. Our dataset describes >46,000 phosphorylation sites across >6600 different proteins, of which ~4500 sites from 2110 proteins show at least a 2-fold increase in response to EGF. We also identified which sites were sensitive to inhibitors of PTPN11, PI3K, MAP2K and RSK using physiological EGF levels. This data was then linked to 15 previously published protein databases to create an interactive web-based resource to explore the relationship between protein phosphorylation, protein characteristics and function.

By correlating the time and dose-dependent changes in signaling pathway protein phosphorylation, we found that RAS-MAPK signaling at low, physiological concentrations of EGF is likely to occur through multiple adaptor pathways that act in parallel. We also found that the level of EGFR phosphorylation was directly proportional to occupancy at all concentrations of EGF and that the kinetics of RAS activation closely paralleled surface bound rather than total EGFR, suggesting that RAS activation is restricted to the cell surface. In addition, we have identified numerous proteins that are likely to be key mediators of the cellular response to MAPK activation. These data have allowed us to extend and refine current models of EGFR-MAPK pathway activation and serve to demonstrate the utility of advanced proteomics technologies for unraveling complex cellular signaling pathways.

## Results

Multiplexed proteomics using isobaric reagents (e.g., tandem mass tags, TMT) can generate precise, large-scale deep datasets across multiple timepoints and conditions (Zecha et al. 2019; Mertins et al. 2018). For use in modeling, however, these timepoints and conditions must be directly relatable to quantifiable physiological outputs that change significantly across those parameters. Thus, to define the most appropriate conditions to evaluate the EGFR-MAPK pathway, we quantified changes in both activated RAS and MAPK across both treatment time and dose of EGF. RAS and MAPK activation were quantified using used enzyme-linked immunoassays (ELISA). Measurement conditions and sample dilutions were optimized to keep all assays within the linear range (*Figure S1 related to Figure 1*).

**Figure 1:**
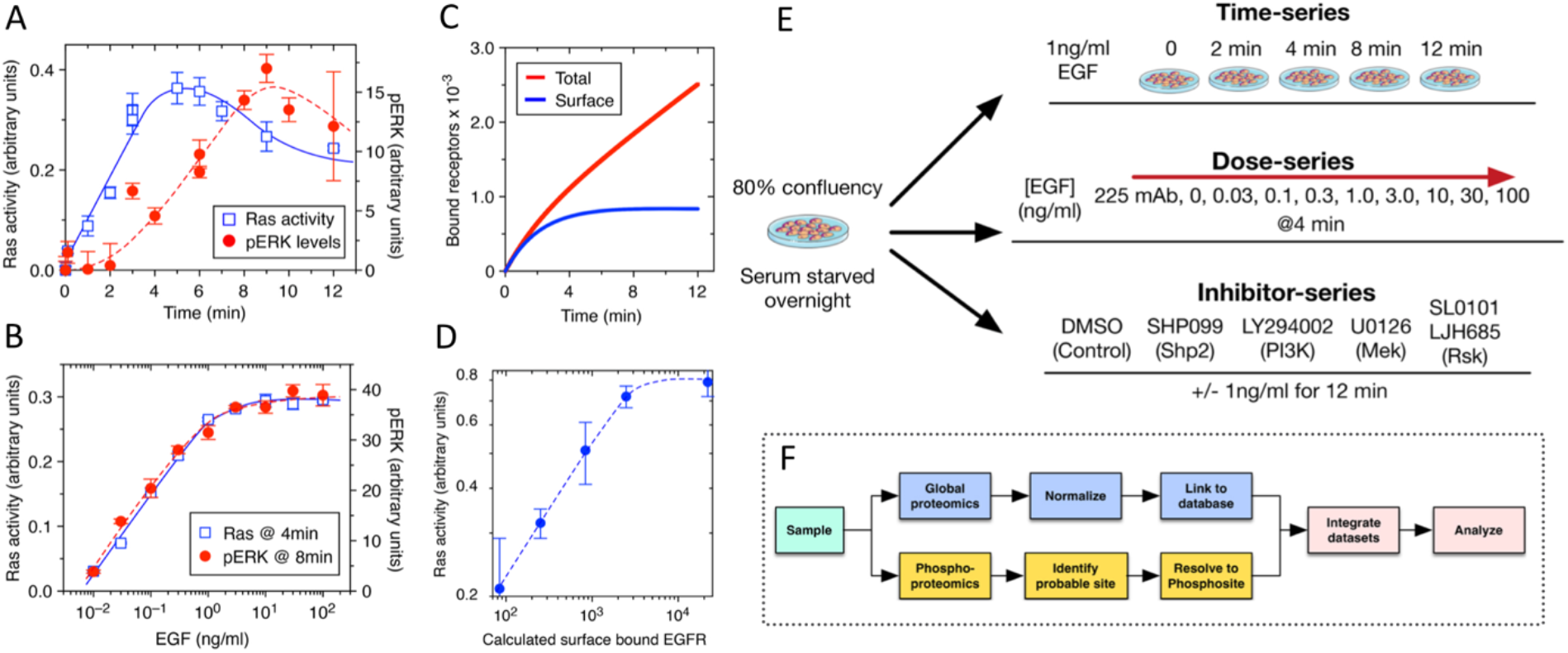
Establishing physiological response range for experiments. 1A: Kinetics of EGF-stimulated RAS and MAPK activation using 1ng/ml EGF. 1B: Dose response of EGF-stimulated Ras and MAPK activation. Samples for each ELISA taken as indicated. 1C: Simulated kinetics of EGF binding and internalization to MCF10A cells. 1ng/ml of EGF treatment was simulated. 1D: Relationship between EGF binding and RAS activation. 1E: Design of sampling experiments. Cells were grown to ~80% confluence and treated with either a range of EGF doses for 4’, or 1ng/ml EGF for 0-12 min. Duplicate samples of each treatment were prepared for multiplexed proteomics. Another set of samples were treated with a variety of inhibitors and then the response to 1ng/ml for 12 min was assessed. 1F: Data analysis and integration workflow. Global proteomics data is normalized to the mean abundance of a panel of cell lines and the phosphoproteomics data resolved to the single site level before integrating with other curated datasets.

In our initial kinetic experiments, cells were stimulated with 1ng/ml of EGF, which has been shown previously to be near optimum for MAPK activation in MCF10A cells (Shi et al. 2016; Gillies et al. 2017). Cells were stimulated for up to 12 minutes, by which time surface binding of EGF has reached a pseudo-steady state and MAPK activation has reached a maximum. As shown in Figure 1A, RAS activity peaks at ~4 minutes whereas MAPK phosphorylation is maximal at 8-9 minutes. This is similar to previous findings as is the ~4-minute delay between maximal RAS and MAPK activation (Joslin et al. 2010; Herrero et al. 2016; Toettcher et al. 2013). We then treated cells with a wide range of EGF concentrations (0.01-100ng/ml) and quantified both activated RAS and pERK at their activation peaks. The identical dose-dependency of the two responses (Figure 1B) demonstrates that the controlling step for MAPK activation is upstream of RAS activation.

In addition to the experimental measurements of RAS and MAPK activation, we generated computer simulations of EGF binding and internalization in these cells. Our models for predicting binding and internalization of EGF have been experimentally validated over decades of research and have proven useful for relating ligand binding to activation kinetics (Resat et al. 2003; Lund et al. 1990). At a concentration of 1ng/ml EGF, surface-bound EGFR reaches a pseudo steady state at ~4 minutes (Figure 1C), similar to the kinetics observed for RAS activation (Figure 1A), whereas total bound EGFR (surface + internalized) continues to accumulate. Interestingly, RAS activation was directly proportional to surface EGFR binding up to an occupancy level of ~2-3,000 receptors per cell (Figure 1D), indicating that below that occupancy level, receptor binding controls RAS activation, but there is an additional limiting step for RAS activation between the EGFR and RAS at higher receptor occupancies.

### Generating the datasets – general approach

The RAS-MAPK functional activity measurements led us to generate three protein phosphorylation profiles: a kinetic profile of between 0-12 minutes, a dose response-series between 0.03 and 100ng/ml, and an inhibitor panel targeting PTPN11, MEK, PI3K and RSK (Figure 1E). Specific sampling points were matched to those used in the functional response studies described above.

We also placed the phosphoproteomics data into the context of the protein expression profile of the cells. This minimizes the likelihood of changes in protein expression being mistaken for changes in enzyme activity. Thus, all samples were evaluated for their global expression of both proteins and phosphoproteins (Figure 1F). Samples were collected in duplicate at the specified time and EGF dose conditions. Typically, 16-plex designs using TMT were used for global proteome and phosphoproteome profiling (Figure 2A). For each type of experiment (e.g., time-series, dose-series, and inhibitor-series), two experimental plexes were used to cover a total of four replicates per condition. The TMT-labeled peptides from all samples for each TMT plex were further subjected to high pH reversed phase fractionation into 12 fractions for global and phosphoproteome profiling (Figure 2A). Quantitative protein abundance data and site-specific phosphorylation data were obtained for each experiment following database searching, processing, and normalization.

**Figure 2:**
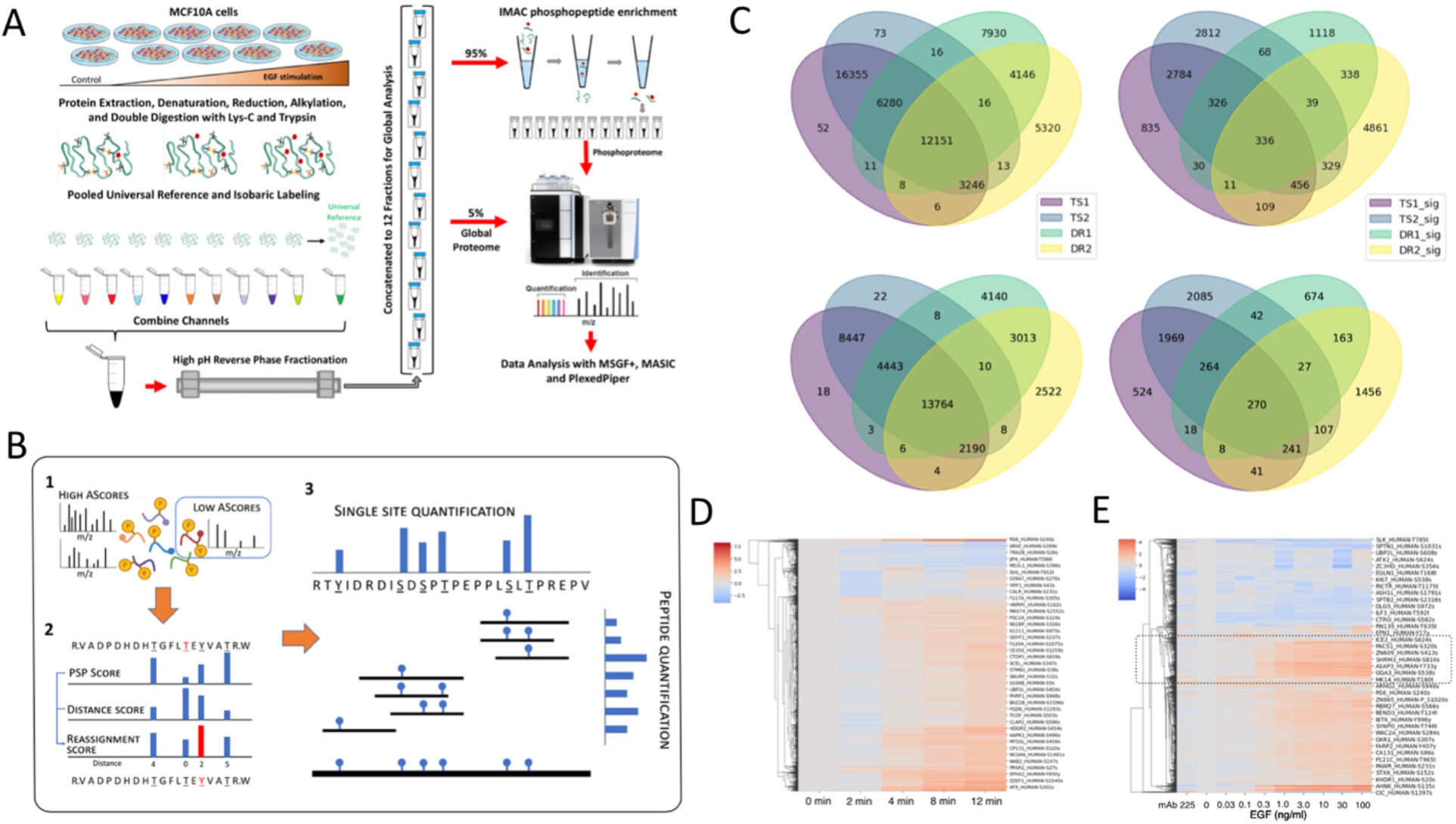
Phosphoproteomics analysis workflow. 2A: Multiplexed quantitative proteomics and phosphoproteomics workflow. 2B: Workflow for resolving phosphorylation data to the single site level. 1) AScore values are combined with 2) a likelihood score based on PhosphoSitePlus data, and 3) Ion intensity values of sites distributed across multiple peptides are then rolled up to a single intensity value. 2C: Impact of remapping site across experiments. Top left: All initially mapped overlapping phosphosites from the two time-series (TS1, TS2) and dose-series (DR1, DR2) experiments. Top right: overlapping phosphosites from sites displaying at least a 2-fold increase in the two time-series (TS1_sig, TS2_sig) and dose-series (DR1_sig, DR2_sig) experiments. Bottom left and bottom right: same data as above following site reassignment and intensity roll-up. 2D: Unsupervised clustering of the remapped significantly changing (2-fold) phosphosites from the time-series experiments. 2E: Unsupervised clustering of the remapped significantly changing (2-fold) phosphosites from the dose-series experiments.

### Linking phosphoproteome data to context

To improve the usability of the phosphoproteomics data, we linked it to a database containing protein-specific data relevant to modeling signaling pathways *(Figure S2 related to Figure 1F)*. These include protein abundance across a wide variety of cell types (Jarnuczak et al. 2021), protein-protein interactions (Huttlin et al. 2021), protein localization (Orre et al. 2019) and functional identity (e.g., adaptor, transcription factors). All MCF10A protein abundance data was normalized to the expression levels of a panel of 132 cell lines (Jarnuczak et al. 2021) and then calibrated to the measured abundance of 46 signaling pathway proteins quantified by targeted proteomics (*Figure S3 related to Figure 1F*). We also linked 17 previously published datasets to our phosphoproteomics data (see Key resources table) to facilitate the analysis of the likely functional impact of specific protein phosphorylation events. This integrated database is available online as Phosphoprotein Explorer (see STAR Methods).

### Resolving phosphopeptides data to single protein sites

A major aim of our study was to provide a reference collection of data that could be linked to previous studies on the relationship between the phosphorylation of specific sites and changes in protein activity. Unfortunately, current approaches for identifying specific modified sites in a phosphopeptide frequently yield ambiguous results. One limitation of tandem mass spectrometry for identifying posttranslational modification (PTM) sites is ambiguity when site-informative fragment ions are missing. Such ambiguity makes it sometimes difficult to link results to sites previously identified in the literature. For example, activated MAPK1 is known to be phosphorylated at residues 185 and 187 (Shi et al. 2015), but our analysis pipeline along with the Ascore algorithm (Beausoleil et al. 2006) would sometimes assign T185 to either T181 or T190. Overall, we found that ~18% of our identified phosphorylation sites were not found in the PhosphositePlus database (Hornbeck et al. 2015). Considering the thousands of samples across hundreds of studies that were used in the construction of this resource, it is highly unlikely that these represent genuine new sites. Our solution was to develop a probabilistic identification scheme that would use both AScore confidence limits (Beausoleil et al. 2006) and prior literature as assembled by the PhosphositePlus database (Figure 2B). This allowed us to reliably map phosphosites between samples as well as improve correspondence of our data to the literature.

An additional problem we encountered was the traditional approach of assigning unique intensity values to multi-phosphorylated phosphopeptides, rather than to their individual phosphorylation sites. If phosphorylation sites are distributed between single and multi-phosphorylated peptides as well as multiple peptides with different peptide lengths due to differences in trypsin cleavage between experiments, all these factors can introduce noise into the data analysis. Distributing intensity values between single and multiple-phosphorylated peptides also makes it difficult to compare results with those from previous site-specific biological literature. We resolved this by remapping the intensity of each parent phosphopeptide to its daughter sites and consolidating sites from multiple phosphopeptides into single intensity values for each specific site (Figure 2B).

Phosphosite remapping and site consolidation significantly reduced the complexity of the dataset. For example, out of 7,580 total proteins detected in our time series study, we identified 38,583 phosphopeptide species from 5,605 proteins. After remapping and reassignment, this dataset was reduced to 28,995 phosphosites from 5,202 proteins, of which >94% could be assigned unambiguously to a single site. Similarly, in the dose series, over 97% of 30,829 phosphopeptides could be resolved to 25,430 single sites. More importantly, remapping improved our ability to compare results across experiments. Out of a total of 55,623 phosphopeptide species observed across the dose and time series experiments, only 22% were observed across all samples (Figure 2C top panels). Following remapping, the total number of unique sites dropped by 30%, but there was a net increase in sites common to all samples, which now constituted 36% of all observed sites (Figure 2C bottom panel). A similar reduction in the complexity of sites changing in response to EGF was also observed (Figure 2C, right panels).

### Properties of the dose and time datasets

Unsupervised clustering of the time-series data yielded a few proteins, such as RPS6 on sites S235-240, that showed strong increase in phosphorylation over time and some that showed a significant decrease, such as ARAF on sites S269-272 (Figure 2D). However, most sites showed a moderate increase in intensity over the 12 min time period. A similar pattern was seen with the dose-series, with a few proteins showing a strong increase or decrease in phosphorylation over increasing EGF dose (Figure 2E). Although a significant group of proteins reached maximum phosphorylation at 1-3 ng/ml EGF (Figure 2E, dotted box), most of the clustered groups showed a gradual increase across all EGF doses (Figure 2E, below dotted box). These data suggest that proteins fall into only a few distinct classes with respect to their response to EGF treatment.

### Validation of the datasets

The proteins in the MAPK pathway that display increased phosphorylation in response to EGF stimulation have been extensively documented over the last several decades and thus provides us with a good basis for evaluating the quality of our data. We first built a “canonical” EGFR-MAPK pathway in MCF10A cells using our previously described perturbation approach (Shi et al. 2016) as well as an extensive literature mining. Our pathway model includes 23 specific proteins of which 20 are phosphorylated (Figure 3A). Since their molecular roles, properties, and sequence in the signaling pathway have been firmly established, these proteins were used to assess the quality and resolution of our phosphoproteomics data.

**Figure 3:**
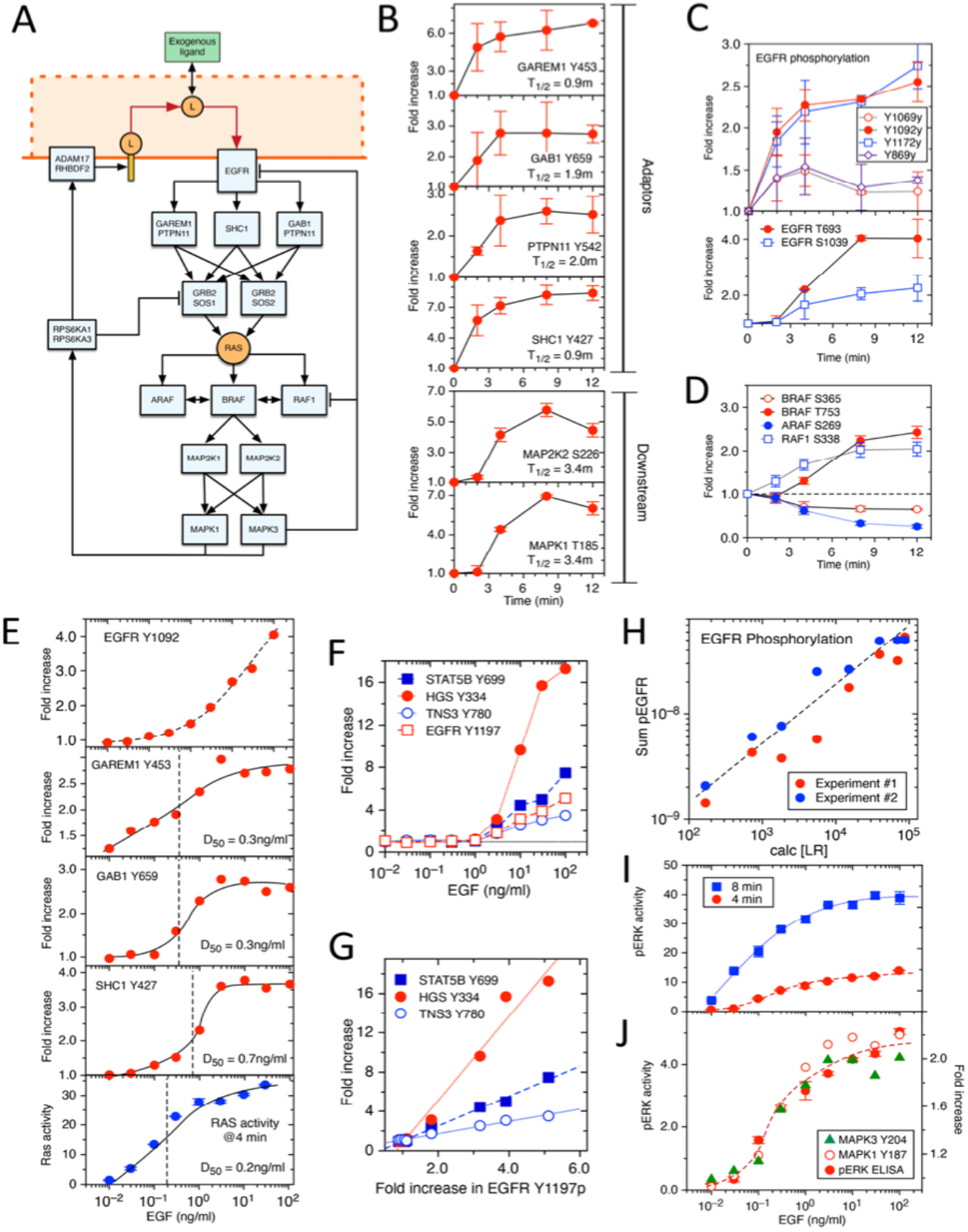
Validation of information flow through pathway. 3A: Map of the EGFR pathway based on previous perturbation experiments. For clarity, only kinase substrates and key intermediate proteins are shown. 3B: Phosphorylation kinetics of key components of the EGFR-MAPK pathway. Adaptors in in the top panels and the MAP2K-MAPK cascade in the bottom ones. Time for half maximum activation estimated by univariate smoothing spline fit to the data. Error bars are SD of duplicate samples. 3C: Phosphorylation kinetics of major regulatory sites on the EGFR. Top panel is the fold increase of four self-phosphorylation sites in the EGFR +/- SD. Bottom panel are negative feedback sites on the EGFR. 3D: Kinetics of both negative and positive phosphorylation of the major Raf proteins. The sites that are decreasing are major inhibitory sites and S338 in RAF1 is an activating site. T753 in BRAF is a negative feedback site. 3E: EGF dose dependency of phosphorylation of key regulatory sites in EGFR-MAPK pathway proteins upstream of Ras. Half maximum dose estimated by – curve fitting. Bottom panel shows the dose-dependency of Ras activation as measured by ELISA. 3F: EGF dose-dependency of self-phosphorylation of EGFR and several downstream direct substrates. 3G: Relationship between self-phosphorylation of the EGFR and downstream substrates. The data in 3F was replotted to show correlation between EGFR self-phosphorylation, indicative of kinase activity, and downstream substrates. Lines are linear regression. All correlation coefficients were >0.99. 3H: Relationship between receptor occupancy and EGFR phosphorylation. The number of occupied receptors at 4 min as a function of dose was calculated as described in Fig 1C and plotted against the sum of the ion intensities of all major self-phosphorylation sites. Shown are the results of two independent experiments. 3I: Time-dependency of MAPK activation. Cells were treated with the indicated dose of EGF for either 4 min (circle) or 8 min (square) before measuring pMAPK levels by ELISA.

### Correspondence to activation kinetics

To evaluate the fidelity of our dataset, we examined the phosphorylation pattern of proteins known to be involved in RAS and MAPK activation. For example, phosphorylation of PTPN11 and GAB1 have been shown to be crucial for RAS activation at physiological (<1ng/ml) EGF concentrations (Dance et al. 2008; Sampaio et al. 2008; Vemulapalli et al. 2021). A comparison between RAS activation kinetics and that of the crucial sites Y259 and Y542 in GAB1 and PTPN11 respectively, show very close similarities (Figure 3B). Similarly, the activation kinetics of MAPK shows a close similarity to the activation phosphorylation at sites S226 on MAP2K2 and T185 on MAPK1. These data show that the phosphorylation patterns we observe correlate well with specific, measurable functions. Phosphorylation of Y427 on SHC1 and Y453 on GAREM1 were also very rapid, in agreement with previous reports (Reddy et al. 2016) and reached maximum at approximately 4 min (Figure 3B), which corresponds to the predicted maximum of EGF binding to surface-associated EGFR (Figure 1C).

Phosphorylation kinetics of the EGFR itself appeared more complex than its downstream substrates. The major self-phosphorylation sites at Y1172 and Y1092 displayed phosphorylation kinetics similar to ligand binding (Figure 3C), with the slow increase after 4 minutes likely reflecting the persistent phosphorylation of internalized receptors (Burke et al. 2001). In contrast, self-phosphorylation sites at Y869 and Y1069 showed a decrease in phosphorylation after 4 min (Figure 3C), which seemed to correspond temporally with an increase in phosphorylation on the negative feedback sites at T693 and S1039 (Figure 3C, lower panel), proposed to be mediated by MAPK and p38 respectively (Takishima et al. 1991; Tanaka et al. 2018). As expected, the kinetics of T693 phosphorylation closely corresponded with the activation of MAPK itself (Figure 3B).

We observed time-dependent phosphorylation of all three RAF isoforms (Figure 3D), although the fold increase for any specific site was modest compared to either upstream or downstream pathway proteins. An increase in phosphorylation of the activation site S338 on RAF1 (Diaz et al. 1997) was paralleled by a decrease in in the inhibitory phosphorylation sites S269 on ARAF (Baljuls et al. 2008) and S365 on BRAF (Hmitou et al. 2007) (Figure 3D) and preceded negative feedback phosphorylation at T753 on BRAF (Ritt et al. 2010) mediated by downstream MAPK. These data show that we can capture the dynamics of both positive and negative phosphorylation events on low abundance proteins in signaling pathways at relevant timescales. Interestingly, although we detected a rapid increase (>2-fold) in the critical tyrosine phosphorylation site Y302 in ARAF (Marais et al. 1997), phosphorylation of the equivalent site Y340/Y341 in RAF1 was not observed (Table S2).

### Dose-response dataset

We evaluated doses of EGF ranging from levels associated with autocrine signaling (<0.1ng/ml; (DeWitt et al. 2001) to those associated with receptor oligomerization (>25ng/ml; (Needham et al. 2016)). A sampling time of 4 min was chosen, which was sufficient for full RAS activation, but only partial MAPK phosphorylation (Figure 1A) to bias the data set towards activating phosphorylation events rather than subsequent negative feedback phosphorylation. In addition, at this time point only ~30% of occupied receptors will have been internalized (Lund et al. 1990). Again, to demonstrate the reliability of the data, we compared the phosphorylation of key activation sites of pathway proteins.

The phosphorylation of the adaptor proteins GAREM1 on its activating site Y453 (Tashiro et al. 2009) as a function of EGF closely paralleled RAS activation, with activating sites on GAB1 and SHC1 showing similar dose dependencies (Figure 3E). All showed a maximal response at ~3ng/ml EGF, which corresponds to the maximum response dose for MAPK activation reported by a variety of investigators (Albeck et al. 2013; Pinilla-Macua et al. 2017; Shi et al. 2016). In contrast, EGFR phosphorylation itself displayed no maximum dose (Figure 3E), suggesting that receptor occupancy had not saturated at the 4 min sampling time at all EGF doses. Interestingly, the dose-response profile of RAS activation most closely resembled that of GAREM1 rather than SHC1 or GAB1 (Figure 3E).

Although EGFR substrates directly involved in MAPK signaling displayed saturation at approximately 3ng/ml, other tyrosine-phosphorylated substrates showed no such effect. As shown in Figure 3F, direct substrates such as HGS, TNS3 and STAT5B showed increasing levels of phosphorylation at EGF doses above 3ng/ml, in parallel with the increasing self-phosphorylation of the EGFR. Indeed, there is a linear relationship between self-phosphorylation of the EGFR at Y1197 or Y1092 and the phosphorylation of non-MAPK pathway substrates (Figure 3G), suggesting that their phosphorylation is directly controlled by EGFR activity. Other phosphoproteins that require high levels of activated EGFR include SH3BP2, TNK2, TNS3 and RBCK1 (Table S3).

To determine the relationship between receptor occupancy and downstream signaling, we converted EGF dose to EGFR occupancy at 4 min. This showed that EGFR self-phosphorylation is directly proportional to EGF binding (Figure 3H), consistent with previous findings (Schooler and Wiley 2000; Shankaran et al. 2012), but inconsistent with reports that activated EGFR can phosphorylate unoccupied receptors (Verveer et al. 2000; Baumdick et al. 2018). This is also consistent with a specific limiting step for RAS activation existing downstream of the EGFR itself (Shi et al. 2016).

MAPK activation showed a similar EGF dose-dependency as RAS with maximum response at about 3ng/ml, at both the maximum activation time of 8 min and at the 4 min time used for the proteomics assay (Figure 3I). This indicates that the dose-limiting factor for MAPK, and presumably RAS activation does not change over the 12min assay time. Significantly, the phosphorylation of MAPK1 and MAPK2 on their activating sites showed an essentially identical profile to the results of the pERK ELISA (Figure 3J), confirming that both assays are in their linear range and that the dynamics of key phosphorylation sites can be correlated with functional assays.

### Inhibitor datasets

Several specific kinase and phosphatase inhibitors targeting EGFR-MAPK pathway proteins have been developed because of the relevance of this pathway to cancer. Specific inhibitors have previously been shown to be an excellent way to dissect the hierarchy of kinase signaling networks, specifically to identify downstream signals that depend on key regulatory proteins (Moritz et al. 2010; Batth et al. 2018). Thus, we determined the impact of inhibiting several key kinases/phosphatases on the pattern of EGF-induced protein phosphorylation, including PTPN11 (Chen et al. 2016), PI3K (Gharbi et al. 2007), MAP2K (Favata et al. 1998) and RSK (Aronchik et al. 2014).

The experiment was run as three 16-plex sets, each with a control and 3 different inhibitors. Cells were pretreated with and without inhibitors, followed by 1ng/ml EGF for 12 minutes. EGF treatment in the absence of inhibitors resulted in at least a 2-fold increase in phosphorylation of between ~700-1300 different resolvable sites in the three experiments (1327, 970, and 696 in sets 1, 2, and 3 respectively). By using a conservative definition of inhibition (see supplementary materials), we found that the PI3K inhibitor LY29400 caused a significant decrease in ~11-22% of the phosphosites whereas the MAP2K inhibitor U0126 significantly reduced 58-90% of the sites (Figure 4A; Tables S4 - S6). Sites reduced with the RSK inhibitors were almost always blocked by MAP2K inhibition as well, which was expected since RSK activation is downstream of MAP2K (Figure 4A). Surprisingly, RSK inhibition significantly decreased ~70% of the sites decreased by MAP2K inhibition alone, suggesting that RSK activity is required for most protein phosphorylation downstream of the MAPK cascade. Sites that were not significantly reduced by any of the inhibitors (~6-35%) included several known direct substrates of the EGFR, such as SHC1 Y427 (Zheng et al. 2013), PTPN11 Y542 (Prahallad et al. 2015) and Eps15 Y849 (Confalonieri et al. 2000). The inhibitors displayed appropriate pathway specificity in that known sites of MAPK phosphorylation, such as EGFR T693 and RAF1 S29 were blocked by the MAP2K, but not the RSK inhibitors whereas the known targets of RSK, such as SOS1 S1134 and ribosomal protein S6 were potently inhibited by the RSK inhibitors (Saha et al. 2012). Similarly, phosphorylation of proteins in the PI3K pathway, such as PRAS40 on S183 (Nascimento et al. 2010), were blocked by the PI3K inhibitor. As expected, PTPN11 Y580 phosphorylation was blocked by the PTPN11 inhibitor (Vemulapalli et al. 2021). Database linkage shows that that MAPK substrates include 13 transcription factors and 12 kinases, whereas RSK substrates include 41 transcription factors and 28 kinases (Figure 4A), confirming that most of the effector functions of the EGFR-MAPK pathway are mediated through RSK. A more complete analysis of the inhibitor data is included in the Supplementary Materials section.

**Figure 4:**
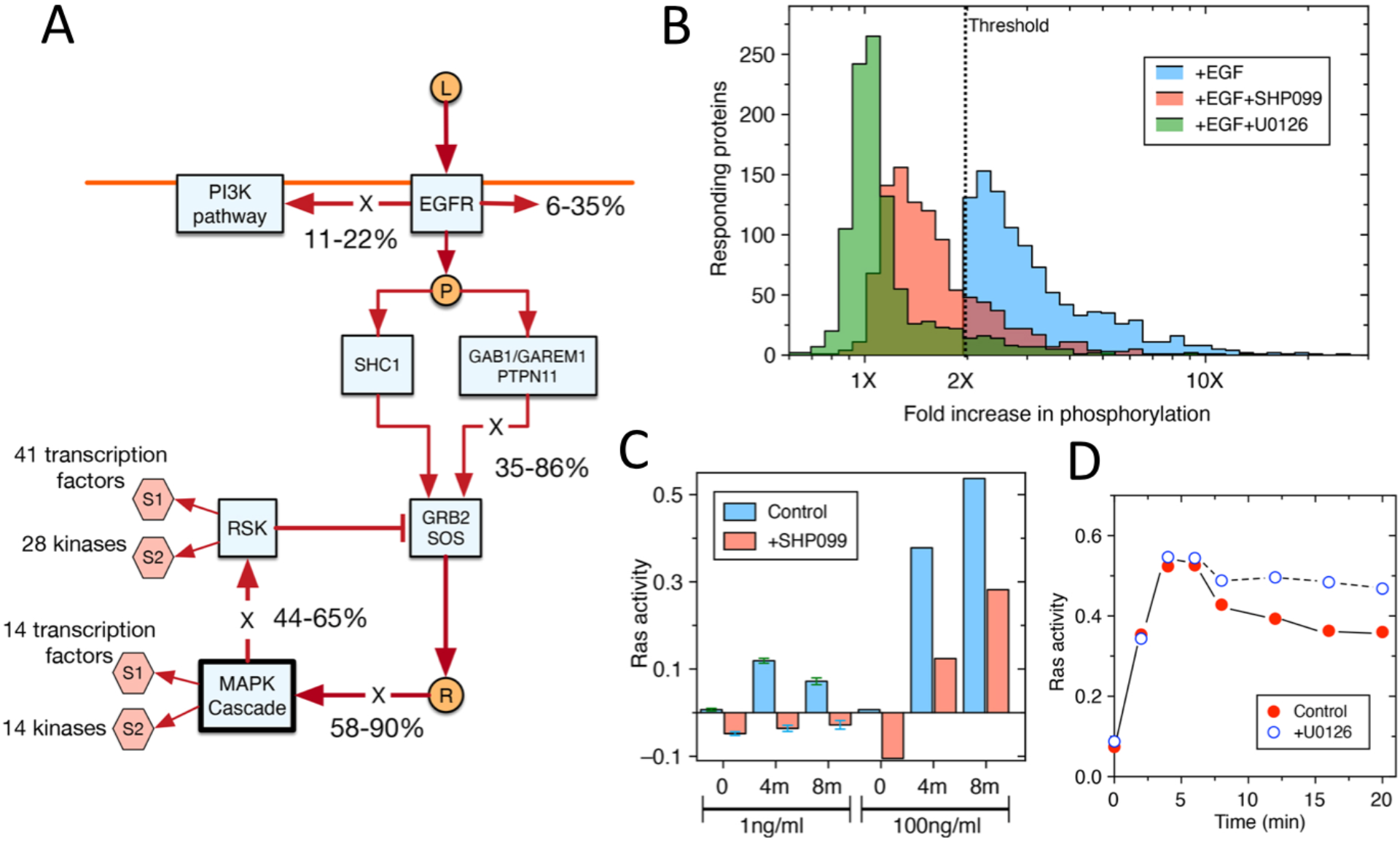
Mapping signal flow through the EGFR-MAPK pathway using selective kinase inhibitors. 4A: Overview of inhibitor results mapped onto the canonical EGFR-MAPK pathway. X marks the know activation steps that are blocked by the inhibitors that were used. Percentages are the range of proteins downstream of the inhibited step that show a significant change in their phosphorylation status. 4B: PTPN11 inhibitor only partially blocks phosphorylation of downstream proteins. Shown is the frequency distribution of proteins display at least a 2-fold increase in phosphorylation (blue) as well as the fold-increase of the same proteins following treatment with SHP099 (red) and U0126 (green). 4C: PTPN11 inhibitor preferentially blocks RAS activation at low levels of EGFR occupancy. Cells were treated with and without 10µM SHP099 for 1 hr before the addition of 1 or 100ng/ml EGF for 4- or 8-min. Ras activity was then measured by ELISA. 4D: MEK inhibitor reduces attenuation of Ras signaling. Cells were treated with and without 10µM U0126 for 1 hr before the addition of 1ng/ml EGF. Cells were assayed for RAS activity at the indicated time points.

We found that in contrast to the MAP2K inhibitor U0126, which was a very strong inhibitor of phosphorylation downstream of the EGFR, the PTPN11 inhibitor SHP099 had a much weaker effect (Figure 4B). From our dose and kinetic data, there appear to be at least three parallel paths that connect EGFR activation to RAS signaling, only two of which are potentially inhibitable by SHP099 (Figure 3A, 4A). Our dose-series data suggest that the PTPN11 partners GAB1 and GAREM1 are phosphorylated at lower EGF concentrations than the alternative adaptor SHC1 (Figure 3E), suggesting that PTPN11 is involved in RAS activation at low EGFR occupancies. To test this idea, we measured RAS activation in the presence and absence of SHP099 at both low (1ng/ml) and high (100ng/ml) EGF concentrations. As shown in Figure 4C, SHP099 reduced basal RAS activity in treated cells and essentially abolished RAS activation by low levels of EGF. However, high levels of EGF partially restored RAS activation, consistent with the idea that PTPN11 preferentially couples low levels of EGFR occupancy to RAS.

We also found that MAP2K inhibition blocked phosphorylation on negative feedback sites, such as SOS1 T1134 (Saha et al. 2012). To determine the impact of these phosphorylation events on signaling, we measured the dynamics of RAS activation in the presences and absence of U0126. As shown in Figure 4D, inhibiting MAP2K greatly reduced the attenuation of RAS activation, at least for the first 20 minutes of pathway activation. This is consistent with previous reports that negative feedback through MAPK/RSK to SOS1 is responsible for the early attenuation of the EGFR-MAPK pathway (Kamioka et al. 2010).

### All phosphosite analysis

Combining data from all three studies (kinetics, dose-response, and inhibitors) based on 6 independent experiments allowed us to quantify a total of 67,771 phosphopeptides from 6642 proteins that could be resolved to 46,804 individual sites. We found that 1287 sites (~3%) could not be unambiguously assigned to a single site and were thus excluded from further analysis. Of the remaining sites, 8,823 (~19%) were observed in all experiments, of which 1283 displayed a >2-fold increase in at least one experiment and 88 displayed a >2-fold increase in all experiments. If our response criterion is expanded to include sites not always observed, then 4,538 sites (~10%) showed a >2-fold increase in at least one experiment.

Because we did not use supraphysiological concentrations of EGF or include phosphatase inhibitors or selectively enrich for pY-containing proteins in our experiments, our data should reflect a relatively unbiased assessment of the MCF10A phosphoproteome. Of all the detected phosphorylation sites, 37,245 were on serine, 7,105 were on threonine and 2,454 were on tyrosine residues, or a S/T/Y phosphorylation ratio of 80:15:5 on 6209, 2730, and 1189 proteins respectively. The ratio between pS and pT (~5.3) is similar to other studies (7.3 – (Olsen et al. 2006) 5.4 – (Sharma et al. 2014), but the number of tyrosine phosphorylation sites we observed is higher than typically reported (Sharma et al. 2014). However, our results are similar to the 74:21:5 distribution reported in a recent reanalysis of phosphoproteomics data from multiple studies from public repositories ((Ochoa et al. 2020).

### Functional characterization of phosphorylation downstream of the EGFR

Our integrated, multi-dimensional dataset allowed us to explore the functional characteristics of protein phosphorylation downstream of the EGFR. As described above, there appears to be a limiting step between EGFR and RAS activation as well as an approximate 4-min lag between RAS and MAPK activation (Figure 1A, Figure 3B). We were interested in whether there were additional time or dose-dependent regulatory steps in the pathway. As shown in Figure 5A, the dose dependency of MAPK and RSK activation are essentially identical, suggesting that the only dose-limiting step in the pathway is between the EGFR and RAS. The kinetics of MAPK and RSK phosphorylation are also essentially identical (Figure 5B), indicating that RSK is activated nearly simultaneous with MAPK. This is also the case with the downstream targets of RSK and MAPK, whose time of half-maximum phosphorylation show a similar distribution (Figure 5C). This is also the case with the phosphorylation of nuclear proteins (Figure 5C), suggesting that the time needed for the transport of activated MAPK into the nucleus does not result in a noticeable lag in substrate phosphorylation.

**Figure 5:**
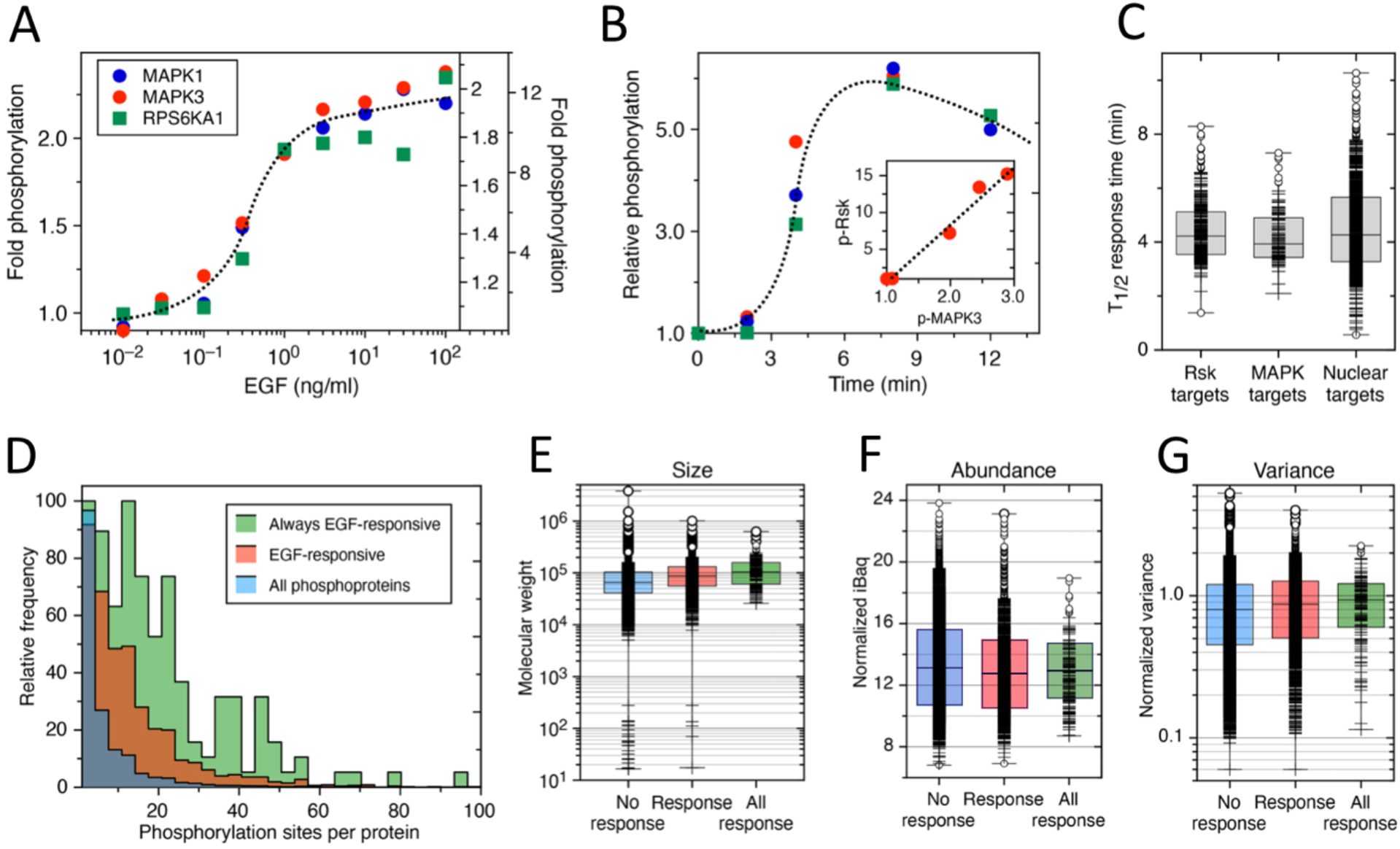
Characteristics of proteins phosphorylated in response to EGF treatment. 5A: EGF dose-dependency of the phosphorylation of the activating site in the kinases MAPK1 (blue circle), MAPK3 (red circle), and RSK1 (green square). The data is scaled to the maximum increase of each site. 5B: Kinetics of phosphorylation of the activating site in the kinases MAPK1, MAPK3, and RSK1 in response to 1ng/ml EGF. The data is scaled to the maximum increase of each site. Same symbols as 5A. 5C: The time for half-maximal phosphorylation of proteins downstream of RSK, downstream of MAPK and classified as nuclear proteins in response to EGF treatment. Phosphorylation is at least 2-fold increase. 5D: Frequency distribution of number of detected phosphorylation sites per protein from MCF10A cells. All phosphoproteins are shown as blue. Proteins that show at least a 2-fold increase in phosphorylation in response to EGF in at least 2 experiments are shown as red. Proteins that displayed at least a 2-fold increase in phosphorylation in all experiments are shown as green. 5E: Molecular weight distribution of proteins classified as in 5D. Box includes the lower and upper quartiles of the data. Close and far outliers as shown as small and large circles respectively. 5F: Abundance distribution of the proteins classified as in 5D. The mean iBaq value of a large panel of cell lines is used as an estimate of protein abundance. iBaq values are logarithmic. Graph description as in 5E. 5G: Mean variance distribution of the proteins classified as in 5D. The variance of protein expression across a large panel of cell lines was normalized to the mean and set as 1. Graph description as in 5E.

We next characterized the proteins that displayed increased phosphorylation in response to EGF treatment. For this effort, we classified all detected phosphoproteins (N=6350) by whether they showed no significant increase in phosphorylation (N=4556), sometimes showed an increase (N=1794) or always showed an increase (N=148). We first looked at the frequency distribution of phosphorylation sites and found that the median number for all phosphoproteins was 4, whereas those that sometimes or always responded to EGF treatment was 9 and 16 respectively (Figure 5D). The size of the proteins was also different between the classes. Non-responding phosphoproteins had a median molecular weight of 85K, the sometime responding protein 109K and the always responding proteins 128K (Figure 5E). However, we saw little difference in either abundance (median iBaq of 13.2, 12.8 and 13.0) or normalized variance (Figure 5F, 5G).

### Inferring signaling network topology from phosphoproteomics data

The depth of our dataset across both time and EGF dose as well as its connection to external sets of protein data allowed us to build several different systems-level maps of the protein phosphorylation network activated by the EGFR. As an example, we first used the inhibitor data to establish the signaling hierarchy and used linked references in the PhosphoSitePlus website to identify functional studies on the biological consequence of specific protein site phosphorylation. This was then used to establish the network topology of positive and negative feedback phosphorylation on all the key proteins in the EGFR-MAPK pathway based on our canonical pathway (Figure 3A). We identified 18 proteins with 41 documented regulatory sites that displayed significant (at least 2-fold) changes in phosphorylation in response to EGF (Figure 6A). Of these, 29 were associated with protein activation and 12 with inhibition (Figure 6B). From these data we reconstructed the regulatory topology of the EGFR-MAPK pathway in MCF10A cells (Figure 6A).

**Figure 6:**
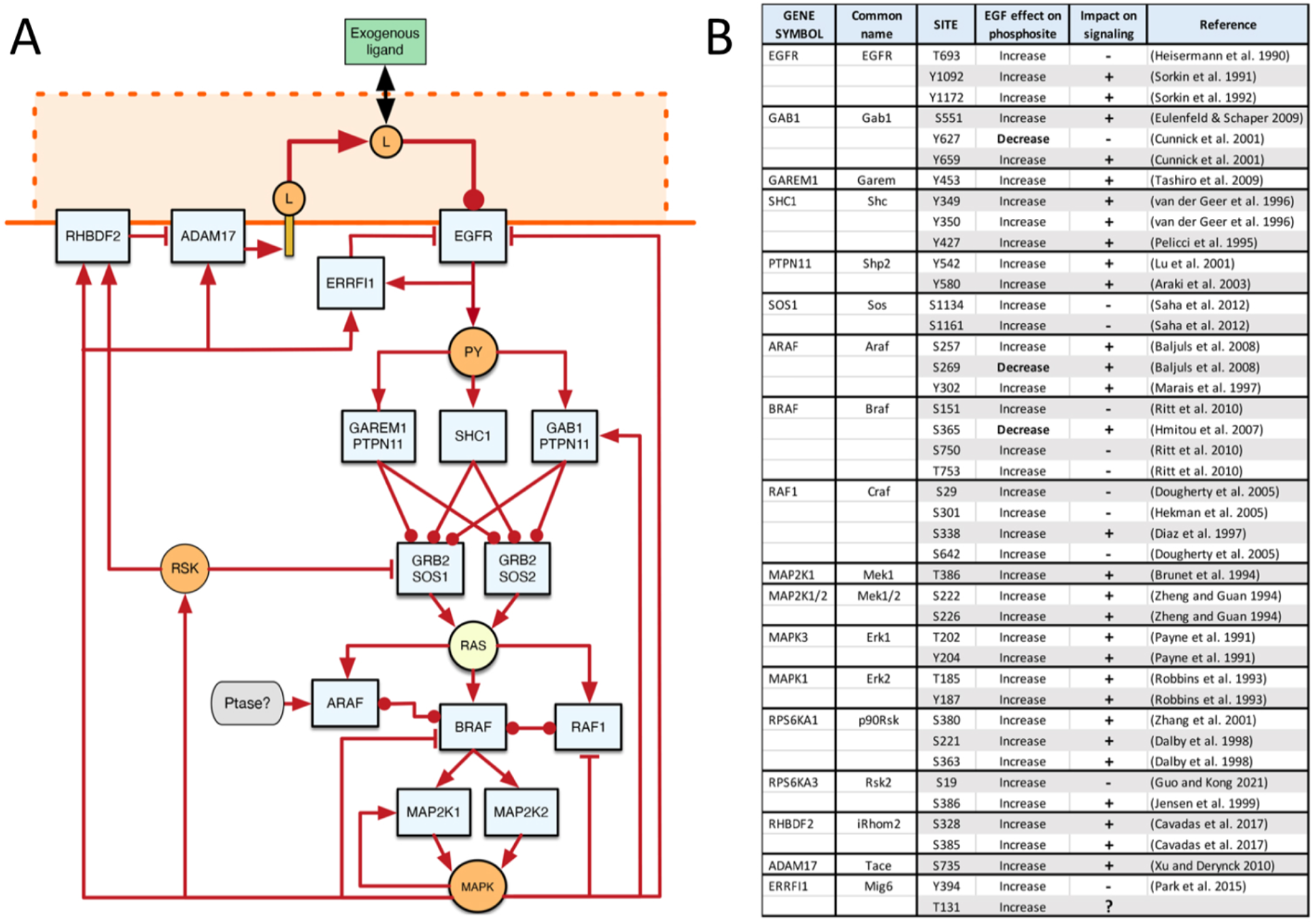
Mapping positive and negative phosphorylation events on the EGFR-MAPK pathway. 6A: Activating events, as defined in the literature, are show by arrows, protein-protein interaction events with round arrows and inhibition events with flat arrows. Multiple activating events are shown as a single arow for clarity. Intermediates that cannot be distinguished from the data are shown as tan circles. 6B: List of specific phosphorylation sites on proteins in the EGFR-MAPK pathway that are known to impact pathway activity. All these sites were modulated at least 2-fold in response to EGF.

We extended our analysis to include the proteins downstream of RAS and MAPK. To be conservative, we restricted our analysis to proteins that displayed at least a 2-fold increase in phosphorylation across all types of experiments. Based on previous studies that suggest that key regulatory proteins are of low abundance (Shi et al. 2016) and show a high number of phosphorylation sites, we further restricted proteins to those that had an iBaq value of <11.5 (~ 10,000 copies per cell) and >0.1 phosphorylation sites per kilobase of molecular weight. This yielded a total of 29 proteins, which included the adaptors of the EGFR, SOS1, and iRhom2 (gene RHBDF2), which regulates ligand shedding and thus autocrine signaling. As shown in Figure 7A, most of these highly phosphorylated EGF responsive (HPER) proteins are downstream of RSK, although some have sites that are independently regulated by MAPK. We found that about half of the regulated phosphorylation sites on HPER proteins have previously been documented to play key roles in the control of specific cellular activities, with the other half being mostly unexplored (Table S7). These proteins are involved in a wide range of cell activities, including intracellular transport, transcriptional regulation, and cytoskeletal organization and migration and the regulated phosphorylation sites show a high median Functional Score (Ochoa et al. 2020).

**Figure 7:**
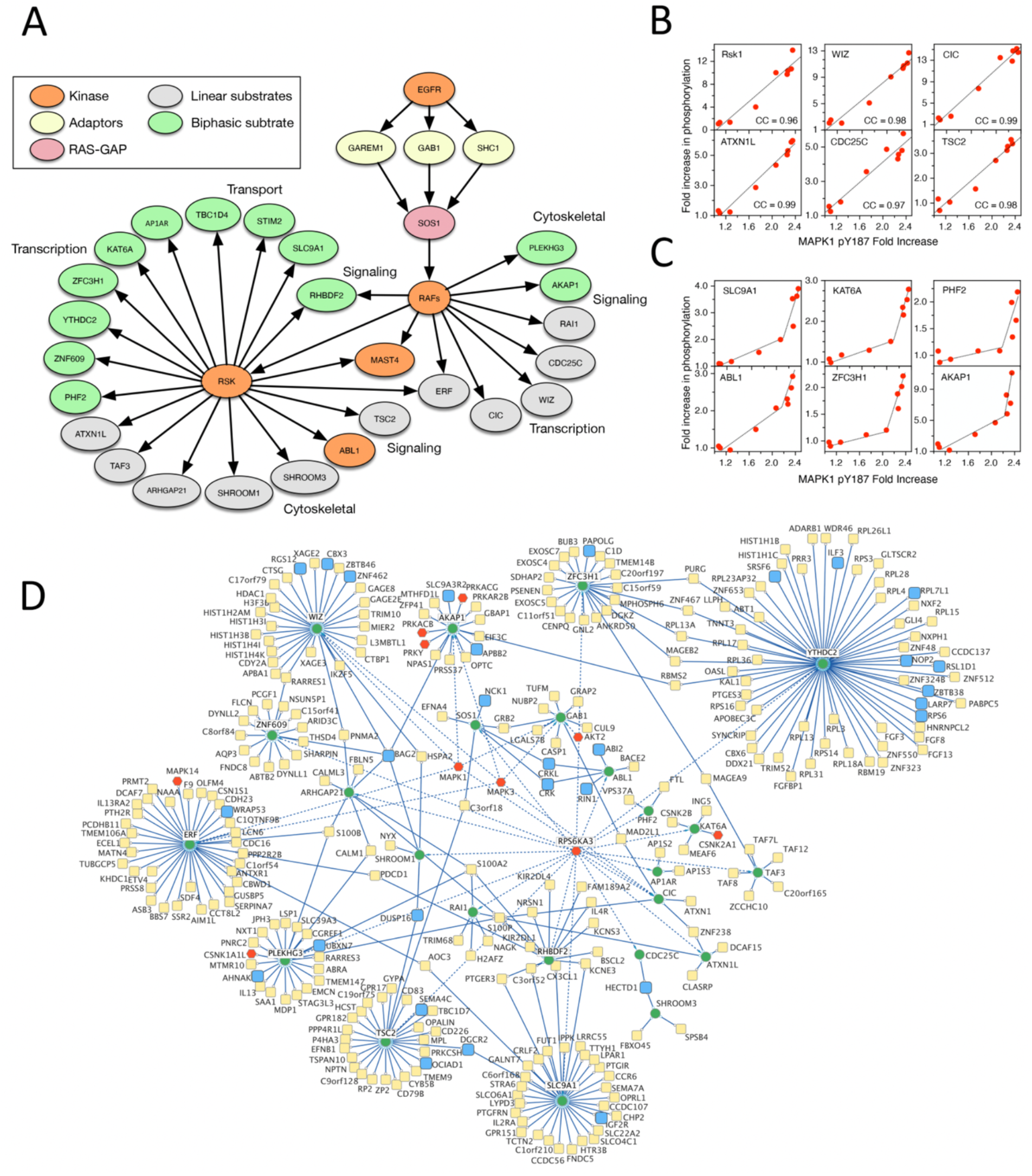
Properties of highly phosphorylated EGF-responsive (HPER) proteins. 7A: Relationship between major HPER proteins as established by kinetics and inhibitor studies. Top of the network is the EGFR (orange). RAFs are shown as a single entity because of our inability to discriminate their downstream specificity. Grey circles are linear responders whereas green are bi-phasic ones. 7B: Linear relationship between MAPK activity and phosphorylation of specific sites in downstream proteins. Representative set of specific phosphorylation sites that show a linear relationship between phosphorylation of the activating site on the major MAPK species in MCF10A cells and specific downstream targets. 7C: Same as 7B, but with selected sites that show a biphasic response to MAPK activation. 7D: Protein interaction network of major HPER proteins. The set of proteins downstream of the Rafs together with their interacting proteins taken from Bioplex are displayed as a Cytoscape network. HPER proteins are green, kinases are red and proteins displaying EGF-stimulated phosphorylation in blue. Inferred relationships are solid lines and kinase dependent relationships are dotted. A very large set of 14-3-3 protein interactions is not shown for clarity.

We demonstrated above that the phosphorylation of RSK and MAPK are directly proportional to RAS activation (Figure 1B, Figure 5A-B). We were curious to see if this relationship held for HPER proteins as well. Thus, we examined the relationship between MAPK activation and downstream HPER protein phosphorylation and found that HPER proteins fell into two, approximately equal classes: those that showed a linear relationship between MAPK activation and their phosphorylation (Figure 7B) and those that showed a linear response up to an EGF dose of 3ng/ml and then a sharp increase thereafter (Figure 7C). As shown in Figure 7A, the linear responders (gray circles) appear in all functional classes of proteins except for transporters.

We next expanded this core network of HPER proteins to include their interaction partners, as documented in the Bioplex database (Huttlin et al. 2021). In Figure 7D, the core HPER network is shown as green nodes, kinases are in red and EGF-stimulated phosphorylated proteins are in blue. Several HPER proteins show extensive protein interaction networks, such as YTHDC2, which is a 3’-5’ RNA helicase thought to regulate mRNA translation and stability (Tanabe et al. 2016). There are eight additional clusters around HPER proteins, but others seem to connect clusters (e.g., ARHGAP21, RHBDF2). Overall, HPER proteins appear to display extensive interactions with each other and other important effector proteins networks.

## Discussion

Here we present a comprehensive atlas of baseline and EGF-induced protein phosphorylation in MCF10A cells across time, dose, and inhibition of multiple kinases. Although several previous studies have identified similar number of total phosphopeptides (Sharma et al. 2014; Batth et al. 2018), those studies used supraphysiological concentrations of growth factors as well as phosphatase inhibitors to increase the number of detected sites. Here, we restricted ourselves to a physiological range of EGF concentrations as well as using a non-transformed cell line expressing a normal complement of EGFR. In parallel with the phosphoproteomics assays, we measured the activation of RAS and MAPK by quantitative ELISA, allowing us to discern relationships between protein phosphorylation and signaling events. Using only 1ng/ml of EGF for a treatment time of <12min, we were able to detect greater than a 2-fold increase in phosphorylation in over 4538 phosphorylation sites on more than 2100 proteins. The depth of this dataset allowed us to evaluate the proposed roles and relationships of many proteins and phosphorylation sites in the activation of the EGFR-MAPK signaling pathway.

Key for this study was the development of a method for reducing the ambiguity of phosphorylation site mapping, allowing the abundance of individual phosphorylation sites to be reliably quantified. Other groups have also identified the issue of ambiguity of phosphorylation site identification and have proposed similar solutions (e.g. (Ochoa et al. 2020; Kalyuzhnyy et al. 2022)). We found that our remapping method, which is based on prior knowledge, reduced noise in our data and improved our ability to integrate data across multiple experiments. Phosphoproteomics data is typically reported as either single or multi-site phosphorylation at the peptide level. Although this can be useful in determining the pattern of phosphorylation on proteins that contain multiple sites, it makes it difficult to integrate the results with the signal transduction literature, which typically looks at the function of individual sites. We have provided our data in both formats to facilitate it use in different types of analyses (Table S8). Our use of highly multiplexed (up to 18-plex) sample tagging together with an improved data analysis pipeline provided us with the most detailed picture of phosphorylation dynamics in the EGFR-MAPK pathway yet reported.

The relatively short time scale (<12min) of our experiments allowed us to measure the initial activation and negative feedback events without the complication caused by new EGF-induced gene expression or protein turnover. We found that between 60-90% of all proteins phosphorylated in response to EGF was dependent on MAPK activation, with most of these depending on RSK activation as well. Relatively few phosphorylation sites depended on PI3K activity, which is consistent with previous studies suggesting that EGF is a poor stimulator of the PI3K pathway in mammary epithelial cells (Rodland et al. 2008).

Because many of the key phosphorylation sites on EGFR-MAPK pathway proteins were observed across all EGF dose, time, and inhibitor treatments, we could evaluate the consistency of our data with previous literature reports. For the most part, we were able to confirm many canonical relationships in the EGFR-MAPK pathway. For example, consistent with previous studies (Zhang et al. 2005; Reddy et al. 2016), we found that activation of the EGFR is essentially coincident with EGF binding (Figure 3C). Phosphorylation of adaptors that bind to the EGFR is also extremely rapid (Figure 3B), as previously reported (Reddy et al. 2016). We also observed an ~4 min delay between RAS activation and MAPK phosphorylation, which appears to be a consistent aspect of MAPK signaling, even when RAS is activated by optogenetics (Toettcher et al. 2013). Our phosphoproteomics data suggest that the delay is at the level of RAF activation since MAP2K phosphorylation shows a similar delay (Fig. 3B). This appears to be the only significant delay in the entire EGFR-MAPK pathway since most substrates downstream of MAPK, even nuclear substrates, appear phosphorylated simultaneous with MAPK activation (Figures 5B-C). We have previously shown that maximal nuclear translocation of MAPK occurs by 5 minutes following EGF addition in human mammary epithelial cells (Shankaran et al. 2009), which indicates that MAPK activation, nuclear translocation and phosphorylation of downstream substrates are so rapid that they are functionally simultaneous. This makes the 4 minutes needed for RAF activation uniquely long and suggests that it could reflect a “kinetic proofreading” process functioning as a low pass filter as previously suggested (Toettcher et al. 2013). Activation of SOS has also been proposed to have a kinetic proofreading step (Iversen et al. 2014), but the time involved (<1min) would not be easily observable with the time resolution of our experiments (~2min).

The delay between phosphorylation of EGFR and MAPK substrates allowed us to detect and temporally discriminate initial versus feedback phosphorylation for multiple proteins. For example the phosphorylation of the activating Y1172 versus feedback T693 on the EGFR could be easily discriminated temporarily (Figure 3C). Interestingly, we observed a decrease in some, but not all the phosphorylation sites on the EGFR at a time that corresponded with phosphorylation of the negative feedback sites (e.g. Y1069; Figure 3C). Although phosphorylation of T693 has been proposed to inhibit the activity of the EGFR (Lan et al. 2019), we have previously shown that it actually alters EGFR substrate specificity (Heisermann et al. 1990). Since the tyrosine phosphorylation sites of the EGFR serve as docking sites of many of its substrates, their selective inhibition could provide a mechanism for changing substrate specificity.

In total, we identified a significant change in the phosphorylation of 29 sites in proteins in the core EGFR-MAPK pathway that have been documented to increase pathway activity and 12 sites documented to decrease pathway activity (Fig. 6). Significantly, many of these proteins function in parallel to each other and negative feedback often only impacts one limb of these parallel pathways. For example, we observed negative feedback phosphorylation on BRAF and RAF1, but not on ARAF, for which we only observed changes on activating sites. We also could not detect a large change in phosphorylation of many of the canonical activation sites on RAF1, such as S259, suggesting either that ARAF is the primary RAF in MCF10A cells, or that many different phosphorylation sites can serve the same functional role as canonical sites. Alternately, the high stoichiometry between RAF1 and the other two RAF isoforms (>2:1) could mean that changes in phosphorylation of only a fraction of the total RAF1 would be sufficient to drive full heterodimerization with BRAF or ARAF (Lavoie and Therrien 2015; Venkatanarayan et al. 2022).

The parallel nature of the EGFR-MAPK pathway is particularly striking in the case of the adaptors that couple activated EGFR to GRB2-SOS. The tyrosine phosphatase PTPN11 appears to be required for binding to GRB2-SOS at low levels of receptor occupancy by forming a complex with either GAREM1 or GAB1 (Figure 4C; also see (Cunnick et al. 2000; Sampaio et al. 2008; Tashiro et al. 2009)). High levels of EGFR occupancy apparently can activate RAS in a PTPN11-independent fashion (Figure 4C), most likely through SHC1 or direct GRB2-SOS binding (Zheng et al. 2013). The parallel and redundant nature of many segments of the EGFR-RAS pathway provides potential mechanisms for “rewiring” of signaling pathways observed when cells develop resistance to cancer drugs targeting the MAPK pathway (Frohlich et al. 2023).

The close correspondence between the phosphorylation pattern of key proteins in the EGFR-MAPK pathway and most previous literature studies provides strong validation of the quality our data. Thus, it should be useful for evaluating several unresolved questions on regulatory mechanisms that control signaling through this pathway. For example, it has been proposed that there are two “classes” of EGFR: high affinity and low affinity receptors that target distinct substrates and give rise to different cellular responses (Krall et al. 2011). At the level of receptor phosphorylation, we see no evidence of multiple classes of EGFR. Instead, self-phosphorylation is directly proportional to EGF binding at all levels of occupancy (Figure 3H). However, we did observe saturation of adaptor phosphorylation and RAS and MAPK activation at relatively low occupancy, but continuously increasing phosphorylation of other EGFR substrates with increasing receptor occupancy. Proteins that are phosphorylated in proportion to all levels of occupancy include HGS, TNS3, STAT5B SH3BP2, TNK2, and RBCK1 (Table S3). These tyrosine-phosphorylated proteins appear to be direct EGFR substrates that do not depend on the adaptors needed for RAS activation. Thus, “high affinity” EGFR could arise from the formation of a limited number of receptor-adaptor complexes involved in RAS activation. “Low affinity” receptors would represent activated EGFR that do not form adaptor complexes because of the limited abundance of adaptor proteins. Significantly, the signaling that occurs at higher levels (>10,000) of EGFR occupancy appears qualitatively different from that mediated by MAPK activation, consistent with a change in substrate preference. We also found that about half of the proteins phosphorylated downstream of MAPK displayed a marked increase in their phosphorylation at high levels of EGFR activation, suggesting that they are targeted by signaling pathways that are activated by both low and high levels of EGFR occupancy. These data suggest that the EGFR pathway can generate qualitatively different cell responses from quantitative differences in receptor occupancy.

The depth and multimodal nature of our dataset also allowed us to define the characteristics of proteins that are consistently phosphorylated in response to activation of the MAPK pathway. To be conservative, we restricted our analyses to proteins whose phosphorylation sites changed at least 2-fold across all experiment types. These 148 proteins are likely only a subset of all downstream kinase targets because of the stringent filtering criteria we applied. However, because these sites were observed across dose, time, and inhibitor series, we can compare their characteristics. In addition, their consistent observation across all conditions means that they are potentially less dependent on cell context.

The most striking aspect of these proteins is their high number of phosphorylation sites. Whereas the median number of sites we observed for all cell phosphoproteins was 4, consistent downstream targets of the MAPK pathway displayed 18 sites (Fig. 5D). The size of these proteins was also larger with a median molecular weight of 128K as compared to 85K for all phosphoproteins. We refer to these as HPER proteins (Highly Phosphorylated EGF Responsive) to reflect their two most prominent characteristics.

To explore the potential regulatory role of HPER proteins, we focused our initial analysis on low-abundance proteins (< ~10,000 copies per cell), based on the previous observation that key regulatory proteins in the EGFR-MAPK pathway tend to be expressed at low levels. This reduced our list from 148 to 29 proteins. This reduced set of low abundance proteins included the key pathway regulators SOS1, GAREM1, GAB1 and iRhom2. The other, non-core pathway HPER proteins were highly enriched for regulators of physiological processes known to be acutely regulated by EGF, such as calcium transport (Chen et al. 1989), endocytosis (Wiley and Cunningham 1982), transcriptional activation (Waters et al. 2012), cytoskeletal dynamics (Felkl et al. 2012), cell migration (Dong et al. 1999) and signal transduction itself (Zielinski et al. 2009). The impact of changes in only about half of the phosphorylation sites have been described in the literature (Table S7), and in those cases phosphorylation on those sites has been shown to have major effects on the activity of the proteins and associated protein complexes. Significantly, the degree of phosphorylation of HPER protein sites was directly proportional to the upstream activity of MAPK itself up to its maximal activation at 1-3ng/ml of EGF. Substrate specificity analysis suggests that approximately 60% of the phosphorylation sites downstream of RSKs could be direct substrates. Although there are likely to be multiple intermediate kinases, such as RSK1 and RSK2, between MAPK and downstream substrates, activation of MAPK is the controlling step.

HPER proteins have several important characteristics. First, although the phosphorylation of specific sites on them are consistently modulated by EGF, these typically constitute a minority of the total sites on these proteins, ranging from 22/50 for CIC to 3/33 for ZNF609 (Table S7). Other kinases have been shown to target several of these proteins, such as AKT, PKA, PKC and JNK, which suggests that their activity can be modulated by kinases from multiple signaling pathways. If so, then the phosphorylation status of these proteins is indicative of the constitutive, baseline kinase activity profile of the cells.

We identify at least 11 HPER proteins that are involved at multiple levels in modulating gene and protein expression. These include KAT6A, PHF2, RAI1 and WIZ, that are involved in histone modification and chromatin remodeling (Rokudai et al. 2013; Baba et al. 2011; Falco et al. 2017; Simon et al. 2015), CIC, ATXN1L, and ERF that normally repress transcription (Ren et al. 2020; Wang et al. 2017; Le Gallic et al. 2004) ZFC3H1 and YTHDC2 that control mRNA retention in the nucleus and translation initiation (Silla et al. 2018; Tanabe et al. 2016) and components of transcription factor complexes, such as TAF3 (Chen et al. 2021). Kinase specificity mapping of the regulated phosphosites strongly suggest that these proteins are direct targets of RSK and MAPK. Thus, the picture that emerges is of MAPK and RSK working together to coordinate the activity of proteins that span the entire hierarchy of regulatory processes that control the transcription and translation of specific proteins.

Some HPER proteins are highly interconnected. By using information from the Bioplex database of protein-protein interactions (Huttlin et al. 2021), we were able to construct a network map of proteins that are likely to be modulated by their phosphorylation state (Fig 7D). Some of the most highly connected proteins, such as YTHDC2, WIZ and ERF are involved in regulating transcription and translation. Other proteins, such as ARHGAP21 and ABL1, have fewer connection but seem to be connected to other HPER proteins through phosphorylated membrane proteins, such as BAG2 or adaptors, such as CRKL (Fig. 7D). Particularly noteworthy is that many of the regulated phosphorylation sites appear to be 14-3-3 protein binding sites (Table S7). 14-3-3 proteins often act as key regulators of protein-protein interactions by sterically hindering or facilitating the formation of specific protein complexes. They are also known to regulate important aspects of EGFR-signaling, such as SOS1-GRB2 binding (Saha et al. 2012), RAF activation (Ritt et al. 2010) and ligand shedding (Cavadas et al. 2017). The high percentage of regulated 14-3-3 sites on HPER proteins suggests that one primary role of EGFR-MAPK signaling is to regulate the assembly of protein complexes that control cell functions by stimulating 14-3-3 binding to key regulatory proteins.

The above is only a small sample of the types of information that can be derived from phosphoproteomics data. Its usefulness is greatly enhanced by using physiologically relevant conditions for generating samples, the use of extensive multiplexing to reduce sample-sample variations, resolving the phosphopeptides data to the single-site level and linking the data to previous, mass-spectrometry-derived protein data resources. All of the data describe here is available as an online, integrated resource to facilitate the exploration of many levels of functional relations. It can also be used as the basis for both validating previous studies and for generating novel hypotheses. It should also be very useful in building a systems-level understanding of how signaling cascades coordinate complex cellular processes and how their dysregulation can lead to disease.

## SUPPLEMENTAL INFORMATION

## ACKNOWLEDGEMENTS

This work was supported by NIH Grants 5U01-CA227544. The experimental work described herein was performed in the Environmental Molecular Sciences Laboratory, Pacific Northwest National Laboratory, a national scientific user facility sponsored by the United States of America Department of Energy under Contract DE-AC05-76RL0 1830.

## AUTHOR CONTRIBUTIONS

H.S.W., H.S. and W.J.Q. conceived and designed the study; H.S.W. and W.J.Q. supervised the work; T.W., C.M.H-B., P.P.D., V.L.P., K.A. and J.A.S. performed experiments, S.F., J.A.S. and H.S.W. analyzed the data; S.F. and H.S.W. wrote software and prepared the integrated database and supplementary tables; H.S.W. wrote the first draft of the manuscript and prepared the figures. All authors reviewed and edited the final manuscript.

## CONTACT FOR REAGENT AND RESOURCE SHARING

Further information and requests for resources and reagents should be directed to and will be fulfilled by the Lead Contact, H. Steven Wiley.

## MATERIALS AVAILABILITY

This study did not generate new unique reagents.

## DATA AND CODE AVAILABILITY

- All mass spectrometry raw data files were deposited in the ProteomeXchange MassIVE database at http://proteomecentral.proteomexchange.org/cgi/GetDataset?ID=PXD044049.
- A FileMaker WebDirect application containing all of the processed data linked to 17 downloaded data resources is available at https://a947342.fmphost.com/fmi/webd/Phosphoprotein_Explorer_V1.
- Python scripts for remapping phosphorylation sites from phosphopeptide data and associated AScores is available from the GitHub site: https://github.com/PNNL-SystemsBiology/EGFR_Phosphoproteomics_Analysis
- R package used at PNNL for processing isobaric labeling (e.g. TMT) proteomics data is available from the GitHub site: https://github.com/PNNL-Comp-Mass-Spec/PlexedPiper

## STAR METHODS

### Key Resources Table

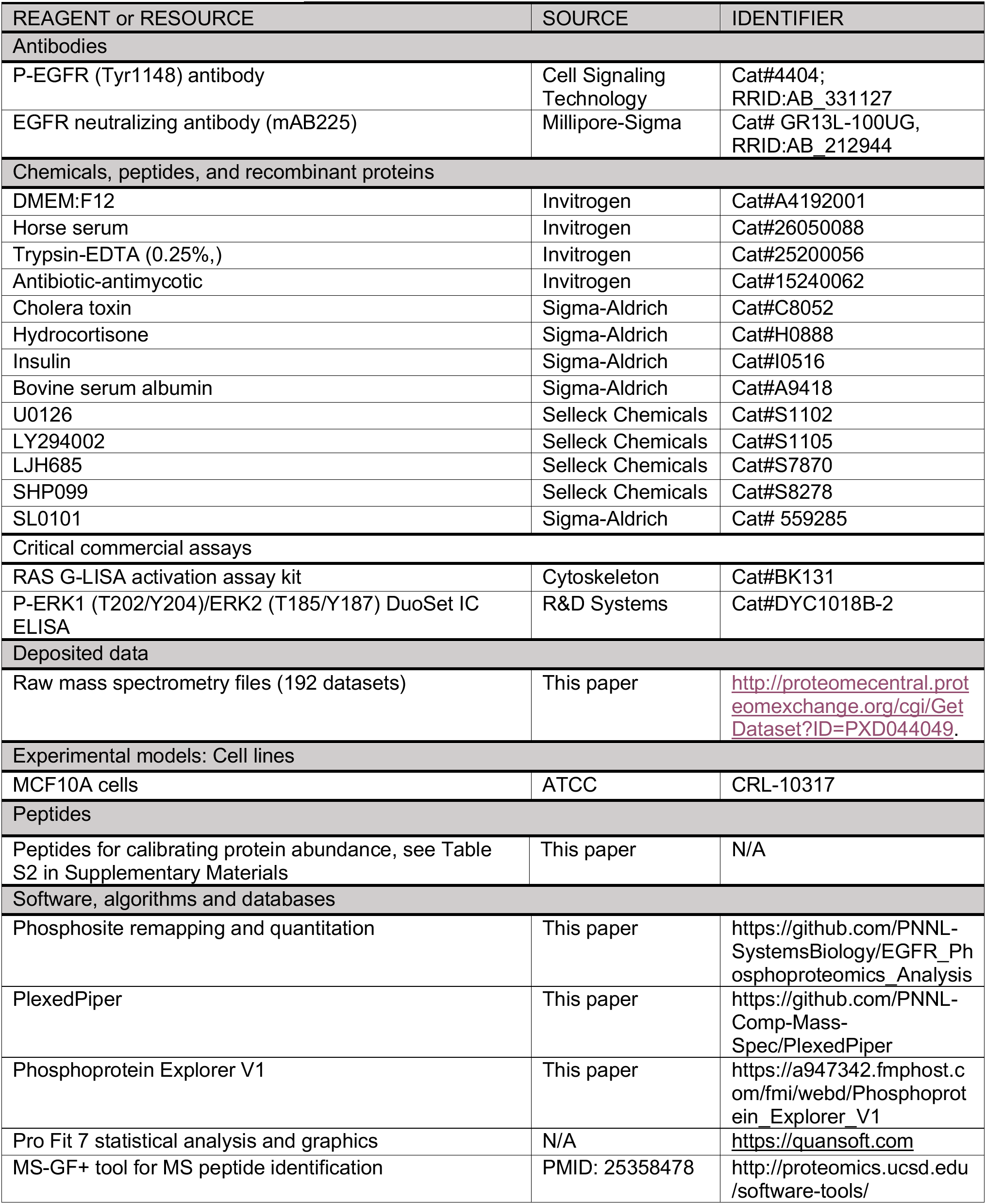
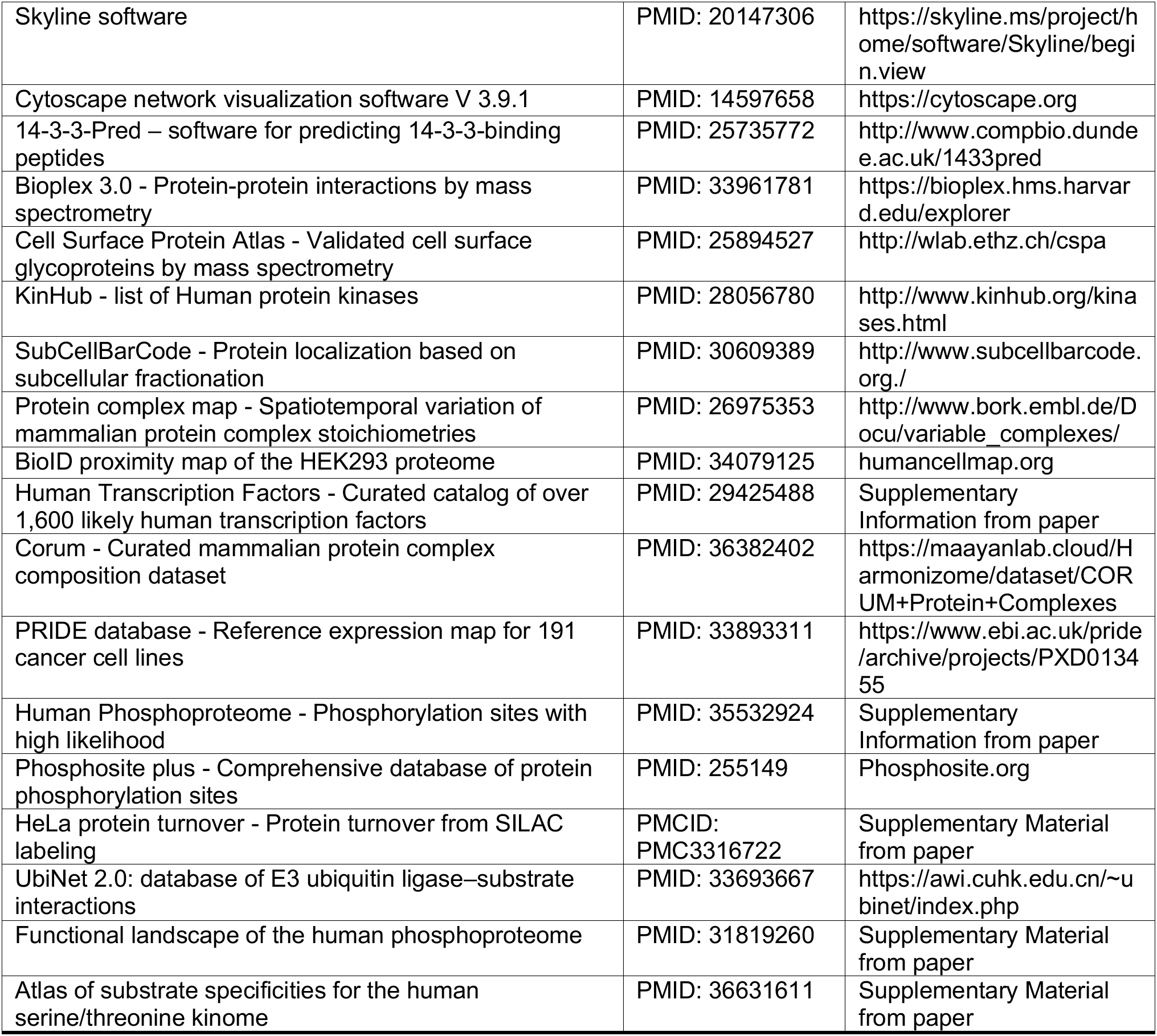

### METHODS DETAILS

#### Cell Culture

MCF10A cells were maintained in growth medium (DMEM:F12, 5% horse serum, 20 ng/ml EGF, 0.5 mg/ml hydrocortisone, 100 ng/ml cholera toxin, 10 ug/ml insulin, 1x antibiotic-antimycotic), subcultured by trypsinization (0.25%) and serum starved in reduced growth factor medium (DMEM:F12, 0.1% bovine serum albumin, 20 ng/ml EGF, 0.5 mg/ml hydrocortisone, 100 ng/ml cholera toxin, 10 ug/ml insulin, 1x antibiotic-antimycotic). For serum starvation, MCF10A cells were washed 2X with HBSS when they reached 40% confluence, followed by addition of reduced growth factor medium for 24 hours (approximately 70% confluence at start of experiments). EGF and selected pharmacons were added directly to the dishes and mixed by gently swirling. Vehicle controls were included in all assays. Typically, cells were cultured in 60 mm dishes using 3 ml medium/dish. Pharmacological inhibitors (15 min pretreatment) were included in select experiments to assess the contribution of target kinases to the measured endpoints. Pharmacons included MEK inhibitor (10 µM U0126); PI3K inhibitor (30 µM LY294002); Shp2 inhibitor (10 µM SHP099); Rsk inhibitors (100 µM SL0101 or 200 µM LJH685); Src inhibitor (5 µM AZD0530).

#### RAS G-LISA Activation Assay Kit

RAS activity was quantified using a G-LISA kit. This kit is based on well-established principles of RAS effector proteins binding to the GTP bound form of RAS as an affinity-based separation method. The kit was run according to manufacturer’s directions. Briefly, lysates were prepared from treated cells by washing cells with ice cold PBS (3X), harvesting monolayers into 70 µl of G-LISA lysis buffer using a rubber policeman, followed by immediate centrifugation at 10,000xg for 1 min at 4°C. The supernatant was rapidly transferred into two Eppendorf tubes. A small aliquot (10 µl) was transferred to one tube for protein determination (BCA), and the second contained the remaining lysate, which was immediately snap frozen in liquid nitrogen and stored at −80°C until experimental analysis.

It is important to note the instability of activated RAS in cell lysates even kept on ice. In our initial characterization of lysates from MCF10A cells, we observed a significant reduction in RAS activity in cell lysates maintained on ice for as little as 40 min before snap freezing in liquid nitrogen. For this reason, it was important to process no more than 3 samples at a time (staggered dosing) so that lysates were only maintained on ice for a few minutes before being snap frozen in liquid nitrogen. To run the G-LISA assay, all reagents were dispensed into the G-LISA plate which was maintained on ice according to the kit directions. The lysates were then rapidly thawed in a 37°C water bath until a small sliver of ice was apparent at which time the tubes were placed back on ice and the appropriate volume of lysate from each tube was pipetted into the G-LISA plate to apply equal protein/well. Preliminary experiments identified 25 µg as an optimal concentration based on the linear range of the assay. Plates were then placed on an orbital shaker for 30 min at 4°C and processed as described in the manufacturer’s directions.

#### ERK ELISA

ERK (i.e. MAPK1 and MAPK3) activation was quantified using a Phospho-MAPK3 (T202/Y204)/MAPK1 (T185/Y187) DuoSet IC ELISA kit according to manufacturer’s directions. Preliminary experiments identified 25 µg as an optimal concentration based on the linear range of the assay (*Figure S1*).

#### Targeted Proteomics Quantification of Protein Abundance

Targeted quantification of protein abundance was performed as we have previously described (Shi et al. 2016). Crude heavy isotope-labeled synthetic peptides were purchased from New England Peptides (now Vivitide) and the top two peptides with the best response were selected to configure the final selection reaction monitoring (SRM) assay. The crude heavy isotope-labeled peptides were spiked into digested peptides of MCF10A cells and analyzed by LC-SRM using the nanoACQUITY UPLC system coupled online to a TSQ Altis triple quadrupole mass spectrometer (Thermo Scientific). For the protein targets that could not be reproducibly detected and quantified by regular LC-SRM, a highly sensitive PRISM (high-Pressure, high-Resolution separations with Intelligent Selection and Multiplexing) method (Shi et al. 2012) was employed prior to LC-SRM analysis to measure their relative abundances. To obtain absolute protein concentration, high-purity light peptides (>95%) were purchased and used to determine the corresponding purity of the spiked-in crude heavy peptides. Based on the peak area ratios of light to heavy peptides, concentrations and copy number of each protein per cell was estimated. All SRM data were analyzed using the Skyline software (MacLean et al. 2010).

#### Protein extraction and digestion

Global and phosphoproteomic profiling was performed using protocols described previously (Hutchinson-Bunch et al. 2021; Mertins et al. 2018; Zecha et al. 2019). Freshly prepared lysis buffer (8M urea, 50 mM Tris-HCl pH 8.0, 75 mM NaCl, 1 mM EDTA, 2 <u>µg</u>/mL aprotinin, 10 <u>µg</u>/mL leupeptin, 1 mM PMSF, 10 mM NaF, 1% phosphatase inhibitor cocktails 2 and 3, and 20 <u>µ</u>M PUGNAc (all reagents from Sigma-Aldrich)) was added to cell pellets, and cells were lysed by vortexing and incubating on a thermomixer set to 4°C and 800 rpm for 30 minutes. Samples were centrifuged for 10 minutes at 4°C and 18,000 rcf to remove cell debris, supernatants were collected, and protein concentration was determined by BCA Assay (ThermoFisher). Protein lysates were denatured with 5 mM dithiothreitol (DTT, Sigma-Aldrich) for 1 hour in a thermomixer set to 37°C and 800 rpm, followed by alkylation with 10 mM iodoacetamide (IAA, Sigma-Aldrich) for 45 minutes in the dark in a thermomixer set to 25°C and 800 rpm. Lysates were diluted 4-fold with 50 mM Tris-HCl pH 8.0 and digested for 2 hours with Lys-C (Wako) at a 1:50 enzyme:substrate ratio in a thermomixer set to 25°C and 800 rpm. Following the initial digest, trypsin (Promega) was added at a 1:50 enzyme:substrate ratio and samples were incubated for 14 hours at 25°C and 800 rpm. Digestions were quenched by adding formic acid (FA) to a final concentration of 1%, samples were centrifuged for 15 minutes at 1500 rcf, and peptides were desalted using C18 solid phase extraction (SPE) cartridges (Waters Sep-Pak). Final peptide concentrations were determined by BCA Assay (ThermoFisher).

#### Tandem Mass Tag (TMT) labeling and basic reversed phase LC fractionation

Peptides were aliquoted for TMT labeling and dried in a vacuum centrifuge, with 250 µg per channel used for TMT16 multiplexes. Peptides were reconstituted in 500 mM HEPES, pH 8.5, to a concentration of 5 µg/µL. TMT reagents were dissolved in anhydrous acetonitrile (ACN) at a concentration of 20 µg/µL and added to peptide samples at a 1:1 peptide:label ratio (by mass). Labeling was allowed to proceed for 1 hour in a thermomixer set to 25°C and 400 rpm, and reactions were quenched with 5% hydroxylamine. Samples from each multiplex were combined, concentrated in a vacuum centrifuge, and cleaned with C18 SPE cartridges. Each combined multiplex sample was fractionated into 96 fractions by high-pH reversed phase separation using a 3.5 µm Agilent Zorbax 300 Extend-C18 column (4.6 mm ID x 250 mm length). Peptides were loaded onto the column in buffer A (4.5 mM ammonium formate, pH 10, in 2% v/v ACN) and eluted off the column using a gradient of buffer B (4.5 mM ammonium formate, pH 10, in 90% v/v ACN) for 96 minutes at a flow rate of 1 mL/minute. After fractionation, samples were concatenated to 12 fractions (Wang et al. 2011).

#### Phosphopeptide enrichment

A small aliquot (5%) of each fraction for each multiplex was removed and vialed at 0.1 µg/µL in 3% ACN + 0.1% FA for LC-MS/MS analysis of global protein abundance. For phosphopeptide enrichment, the remaining 95% of each fraction was dried by vacuum centrifuge and reconstituted to 0.5 µg/µL with 80% ACN + 0.1% TFA. Fe^3+^-NTA-agarose beads were freshly prepared using Ni-NTA-agarose beads (Qiagen), and 40 µL of bead suspension was added to each fraction and incubated at room temperature for 30 minutes on a thermomixer set to 800 rpm. After incubation, beads were washed with 100 µL of 80% ACN+0.1% TFA and 50 µL of 1% FA to remove nonspecific binding. Phosphopeptides were eluted off the beads with 210 µL (3 x 70 µL elutions) of 500 mM K_2_HPO_4_, pH 7.0, directly onto C18 stage tips and then eluted from C18 material with 60 µL of 50% ACN +0.1% FA. Phosphopeptide samples were dried in a vacuum centrifuge and reconstituted with 12 µL of 3% ACN +0.1% FA immediately prior to LC-MS/MS analysis.

#### Proteomics data processing

The obtained MS/MS spectra were searched against the UniProt human database (downloaded November 5^th^ 2019) using the MS-GF+ tool (Kim and Pevzner 2014) for peptide sequence identification. Carbamidomethylation on cysteine residues and TMT modifications on lysine and peptide N-termini were set as fixed modifications, with methionine oxidation set as a dynamic modification. For phosphoproteomics datasets, phosphorylation on serine, threonine, and tyrosine residues was set as an additional dynamic modification. Localization of phosphorylation sites was performed using the AScore algorithm (Beausoleil et al. 2006). A target-decoy approach was used to control false discovery rates of peptide and protein identifications to 1%. For identified peptides, TMT reporter ion intensities were extracted by the MASIC tool (Monroe et al. 2008). Data from all fractions were aggregated for each multiplex based on common peptides or proteins (global proteomics) or phosphopeptides (phosphoproteomics) and median-centered using in-house developed R package (https://github.com/PNNL-Comp-Mass-Spec/PlexedPiper).

Data from individual samples were median-centered across each multiplex before comparisons were made. We found that this was a more reliable way to normalize the phosphoproteomics data than using the global data because variations between individual sample processing was more significant than the changes in phosphorylation state induced by the experimental treatments (*Figure S3*).

#### Phosphorylation reassignment

The procedure for reassignment and quantification of sites on phosphopeptides:

1. For each identified phosphopeptide with a high AScore (Beausoleil et al. 2006) (> 60) their site was assigned an experiment score (*S_E_*) of 1. All lower scoring phosphopeptides were selected for reassignment.
2. For each candidate peptide for reassignment, we identified all possible phosphorylation sites and assigned each two likelihood scores using the PhosphoSitePlus evidence: mass-spectrometry (MS) based and low-throughput (from the literature) based. These two scores are calculated from the number of observations at each site in that peptide divided by the number of total observed sites in that peptide. We normalize these two types of observations separately as MS score *S_M_* and literature score *S_L_*.
3. Based on the position of the possible phosphorylation site relative to originally assigned site(s), we calculate a distance score *D* for each candidate site with *D* = AScore/(*d* + 1), where *d* is the smallest distance between the candidate site with originally assigned site(s). This is to account for the limited information contained in the fragments in the cases of non-zero AScores. We then normalize the distance scores by dividing the sum of *D* values from all candidate sites within the peptide, which results in *S_D_* scores.
4. We sum up these scores with user defined heuristic weights to get the final scores *S_F_* for ranking the likelihood of all candidate sites, using the equation:

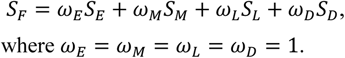
5. We then take the top *n* sites with the highest *S_F_* scores and assign them as the phosphorylated site(s) of the peptide, where *n* is the number of phosphorylation sites that were originally assigned to the peptide.

After reassignments, we sum the intensity values of all phosphopeptides containing a specific site to yield the total intensity value of that site.

#### Quantifying impact of inhibitors on phosphorylation sites

Because of the sample-sample variability we observed in the phosphoproteomics data, there is significant potential for false positive and false negative assignments for both EGF-responsive and inhibitor-responsive sites. In addition, the inhibitors are not 100% efficient in their target activity and off-target activities will always be present to some extent. To be conservative in our assignment of EGF-responsive sites that are inhibited by specific treatments, we used the following criteria for each detected phosphorylation site:

1. A response to EGF by at least a 2-fold increase in average ion intensity (AII) of a specific site relative to the control (-EGF) value in inhibitor treated and untreated cells.
2. A difference between the AII of a site between inhibitor treated and untreated samples > sum of the standard deviations in fold-response.
3. The inhibitor must reduce the magnitude of the EGF response by at least 50%

We calculated the standard deviation of the fold-change by using the fold change in all the paired replicates (+E1/C1, +E2/C1, +E1/C2, +E2/C2) to generate 4 estimated fold-changes and taking the standard deviation of these values. Mean values were calculated from the average of the four ratios.

#### Calculation of half-max response time *T*_1/2_ and half-max response dose *D*_1/2_

For *T*_1/2_, we calculate the Log2 fold changes of each phosphorylation site at 2 min, 4 min, 8 min, and 12 min relative to 0 min. Then we use the univariate smoothing spline method to interpolate the Log2 fold changes corresponding time changes. The roots of the interpolated functions are calculated for finding the maximums and minimums. We take the maximum or minimum that has the largest absolute value as the response max. Then we use the interpolation function to locate the max response time *T*_Max_, as well as the half-max response time *T*_1/2_ between 0 and *T*_Max_. Similarly, for *D*_1/2_, we calculate the Log2 fold changes of each phosphorylation site treated with 0.03 ng/ml, 0.1 ng/ml, 0.3 ng/ml, 1.0 ng/ml, 3.0 ng/ml, 10 ng/ml, 30 ng/ml, 100 ng/ml EGF as well as mAb 225 relative to the one treated with DMSO (i.e., 0 ng/ml EGF). Then we use the same interpolation method and root function to locate the max response and half-max response as well as the doses corresponding to them.

#### Estimating EGFR occupancy

The level of binding of EGF to its receptor at any given time point was calculated by numeric integration of the rate equations describing the forward and reverse rate constants as well as receptor internalization (Knauer et al. 1984). These equations accurately describe the dynamics of cells interacting with the EGFR at short time intervals (<15 min; (DeWitt et al. 2001) Rate constants that were used were: k_a_ = 1.2 x 10^6^ M^-1^s^-1^; k_d_ = 3.67 x 10^-2^s^-1^; k_e_ = 4.0 x 10^-3^s^-1^; k_t_ = 1.17 x 10^-3^s^-1^. The value of V_r_ (292) was adjusted to yield the initial number of EGFR derived from targeted proteomics measurements (2.5 x 10^5^). The simulated volume was 2ml and cell number was 1.2 x 10^6^. Calculated occupied receptors included both receptors at the cell surface as well as internalized (Lund et al. 1990).

#### Integrating phosphoproteomics data with protein information

To improve the usability of the phosphoproteomics data, we linked it to a database containing protein-specific data relevant to modeling signaling pathways (*Figure S2*). These include protein abundance across a wide variety of cell types, protein-protein interactions, protein localization and functional identity (e.g., adaptor, transcription factor, etc.). In all, we normalized and linked 17 previously published datasets to our phosphoproteomics data, which greatly facilitated identification of the likely functional impact of specific protein phosphorylation events (listed in Key Resource Table).

Most data were linked through UniProt or Ensembl IDs, depending on original data tables. A cross-reference table was constructed to allow the addition of new data with different types of identifiers. Conversions from UniProt IDs to GeneIDs were most efficiently performed using David v6.8 (Huang da et al. 2009). IPI codes were translated into GeneIDs using a custom database. Gene Symbols and Ensembl IDs were directly generated from the NLM Gene database.

All protein abundance data was normalized using the results from the study of Jarnuczak *et al*. (Jarnuczak et al. 2021), which reports the average characteristics of 340 proteomics samples from over 100 different cell lines. The iBAQ abundance values of each sample were normalized using their advanced algorithm, providing a good benchmark against which to normalize new proteomics datasets. We report the average, normalized log2 iBAQ value for each protein with the average log2 iBaq value of all proteins from all samples across all cell lines being 13.6.

We mean-centered our data to the pooled data to generate normalized iBAQ values for each protein we detect in MCF10A cells. These values and those of from pooled dataset are included as well as the normalized variance values (variance-mean ratios) for each protein calculated from the pooled datasets. Although the inferred data values from the pooled datasets were included for the abundance normalization calculations, they were removed for both average protein abundance estimation and variance calculations. The fraction of cells that express the indicated protein is also provided.

iBaq values were converted to estimated copies per cell by first quantifying the levels of 45 signaling proteins in MCF10A cells by targeted proteomics as described above using purified (>95%) isotopically labeled peptides as internal standards (see Key Resource Table). Protein abundance values ranged from 1600 to 730,000 copies per cell in this assay. Regression analysis of log2 copies per cell versus iBaq values yielded a correlation coefficient of 0.61 and a significance of correlation of 0.03% with a slope of 0.43 and Y-intercept of 9 (*Figure S3*). The correlation we observe between iBaq and SRM-derived values are similar to what has previously been reported (Jarnuczak et al. 2021) and should only be considered a general estimate in lieu of direct measurements.

The data from the sources listed in Table S2 were imported into a FileMaker Pro relational database (version 19.6) and linked through the indicated identifiers. This allowed for searches based on either the properties of the phosphoproteomics data or parameters contained in any of the linked datasets. For example, a search for sites that respond to EGF, and the identified protein kinases returns a list of 112 kinases that are phosphorylated in response to EGF treatment. This database is available online as <u>Phosphoprotein Explorer</u> and as a freestanding application file upon request.

