## Supplemental Table 1 for "A Phosphoproteomics Data Resource for Systems-level Modeling of Kinase Signaling Networks"

### Table S1 Targeted peptides used for calibration of iBaq values for protein abundance

| Peptides | Source |
|---|---|
| ADAM17 : NIYLNSGLTSTK | Thermo Fisher Scientific |
| AKT1 : FYGAEIVSALDYLHSEK | Thermo Fisher Scientific |
| APC : SESEDLQQVIASVLR | Vivitide |
| ARAF : TQHCDPEHFPFPAPANAPLQR | Thermo Fisher Scientific |
| AXIN1 : SDIYLEYTR | Vivitide |
| AXL : GQTPYPGVENSEIYDYLR | Vivitide |
| BRAF : LLFQGFR | Vivitide |
| CBL : GTEPIVVDPFDPR | Thermo Fisher Scientific |
| CSNK1A1 : ILQGGVGIPHIR | Vivitide |
| CTNNB1 : NEGVATYAAAVLFR | Vivitide |
| EGFR : LTQLGTFEDHFLSLQR | Thermo Fisher Scientific |
| ERRFI1 : NSPSLFPCAPLCER | Thermo Fisher Scientific |
| GAB1 : LTGDPDVLEYYK | Thermo Fisher Scientific |
| GRB2 : ATADDELSFK | Thermo Fisher Scientific |
| GSK3B : LLEYTPTAR | Vivitide |
| H/K/NRAS : LVVVGAGGVGK | Thermo Fisher Scientific |
| HRAS : SFEDIHQYR | Thermo Fisher Scientific |
| IGF1R : LGCSASNFVFAR | Vivitide |
| INPPL1 : LLLDTLQLSK | Vivitide |
| iRHOM1 : GTADWFGVSK | Vivitide |
| iRHOM2 : DCSETLATFVK | Vivitide |
| IRS1 : LGPAPPGAASICRPTR | Vivitide |
| KRAS : SFEDIHHYR | Thermo Fisher Scientific |
| MAP2K1 : ISELGAGNGGVVFK | Thermo Fisher Scientific |
| MAP2K1/2 : LIHLEIKPAIR | Thermo Fisher Scientific |
| MAP2K2 : ISELGAGNGGVVTK | Thermo Fisher Scientific |
| MAPK1 : VADPDHDHTGFLTEYVATR | Thermo Fisher Scientific |
| MAPK3 : IADPEHDHTGFLTEYVATR | Thermo Fisher Scientific |
| MET : VADFGLAR | Vivitide |
| NRAS : SFADINLYR | Thermo Fisher Scientific |
| PDPK1 : QLLLTEGPHLYYVDPVNK | Thermo Fisher Scientific |
| PEBP1 : LYTLVLTDPDAPSR | Thermo Fisher Scientific |
| PIK3CA : FGLLLESYCR | Vivitide |
| PIK3R1 : FSAASSDNTENLIK | Thermo Fisher Scientific |
| PIK3R2 : ALGATFGPLLLR | Thermo Fisher Scientific |
| PLCG1 : IGTAEPDYGALYEGR | Vivitide |
| PTPN11 : SNPGDFTLSVR | Thermo Fisher Scientific |
| PTPRE : TGTFIALSNILER | Thermo Fisher Scientific |
| RAF1 : TISNGFGFK | Thermo Fisher Scientific |
| RASA1 : LLAITELLQQK | Thermo Fisher Scientific |
| SHB : GAEHLSLLYPVAVR | Vivitide |
| SHC1 : EAISLVCEAVPGAK | Thermo Fisher Scientific |
| SHOC2 : LDSLTTLYLR | Thermo Fisher Scientific |
| SOS1 : LPSADVYR | Thermo Fisher Scientific |
| SOS1 : EYIQPVQLR | Thermo Fisher Scientific |
| SPRY4 : LQHPLTILPIDQVK | Thermo Fisher Scientific |
| STAT3 : GLSIEQLTTLAEK | Vivitide |
