## Supplemental Table S7 for "A Phosphoproteomics Data Resource for Systems-level Modeling of Kinase Signaling Networks"

| Gene Symbol | Uniprot ID | # Psites | # EGF-reg sites | Sites/KB MW | iBaq | Variance | Regulated site | Functional score | Fold change @1ng | T Max | Dose response | Upstream kinase | Cannonical kinase | Reference | 14-3-3 Prediction | Protein function | Impact of site phosphorylation |
|---|---|---|---|---|---|---|---|---|---|---|---|---|---|---|---|---|---|
| TBC1D4 | O60343 | 22 | 4 | 0.15 | 10.87 | 0.10 | S588 | 0.49 | 2.4 | 5.8 | Bi-phasic | RSK | AKT | PMID: 12637568 | + | Regulates translocation of glucose transporters | Induces GLUT4 translocation through inactivation of its Rab GAP function. |
| STIM2 | Q9P246 | 14 | 3 | 0.17 | 10.03 | 0.83 | S699 | 0.47 | 5.3 | 8.0 | Bi-phasic | RSK | | PMID: 29783744 | ++ | Regulates calcium transport and distribution in cells | Not known |
| AP1AR | Q63HQ0 | 6 | 2 | 0.17 | 10.71 | | S175 | 0.38 | 3.8 | 7.2 | Bi-phasic | RSK | | PMID: 19706427 | - | Regulator of the AP-1 adaptor complex involved in vesicular trafficking. | Not known |
| SLC9A1 | P19634 | 10 | 4 | 0.13 | 11.34 | 1.34 | S703 | 0.75 | 12.7 | 6.1 | Bi-phasic | RSK | RSK | PMID: 25578862 | +++ | Sodium/hydrogen exchanger 1. Involved in pH regulation to eliminate acids generated by active metabolism or to counte | Activates Na+/H+ transporter activity, enhancing cell migration |
| KAT6A | Q92794 | 27 | 5 | 0.10 | 8.71 | 1.10 | S448 | 0.26 | 3.3 | 11.8 | Bi-phasic | RSK | AKT | PMID: 23431171 | - | Co-activator of transcription by acetylation of histones | Not known |
| PHF2 | O75151 | 20 | 5 | 0.16 | 9.78 | 1.11 | S853 | 0.18 | 2.2 | 10.0 | Bi-phasic | RSK | PKA | PMID: 21532585 | + | Histone lysine demethylase | Activates histone lysine demethylase activity, thus activating transcription |
| ZNF609 | O15014 | 33 | 3 | 0.22 | 10.69 | 1.38 | S413 | 0.56 | 5.0 | >12 | Bi-phasic | RSK | | PMID: 23076336 | + | Transcription factor, predicted to enable promoter-specific chromatin binding activity. | Not known |
| YTHDC2 | Q9H6S0 | 17 | 3 | 0.11 | 11.21 | 0.85 | S1202 | 0.39 | 2.0 | >12 | Bi-phasic | RSK | | PMID: 26996300 | - | Member of a family of ATP-dependent RNA helicases that promotes translation initiation by recognizing and binding N6 | Not known |
| ZFC3H1 | O60293 | 23 | 9 | 0.10 | 10.21 | 0.93 | S352 | 0.42 | 4.9 | 11.1 | Bi-phasic | RSK | | PMID: 29768216 | +++ | Acts as a central nuclear pA+ RNA retention factor, counteracting nuclear export activity, generally targeting longer and | Not known |
| WIZ | O95785 | 21 | 8 | 0.11 | 11.36 | 0.85 | S982,S983 | 0.19/0.31 | 2.7-10.3 | 6.2-8.4 | Linear | MAPK | | PMID: 26338712 | - | Regulates histone methyltransferase assembly and activity, thus regulating transcription | Not known |
| ERF | P50548 | 17 | 8 | 0.31 | 9.32 | 1.23 | S21,S531 | 0.65/0.67 | 2-6.5 | 10-12 | Linear | MAPK/RSK | MAPK | PMID: 14729966 | +++ | Potent transcriptional repressor that binds to the H1 element of the Ets2 promoter. | Induces export from the nucleus, thus activating transcription and cell cycle progression |
| RAI1 | Q7Z5J4 | 36 | 6 | 0.19 | 10.86 | 0.53 | T1068 | 0.41 | 4.3 | 6.5 | Linear | MAPK | | PMID: 29138588 | - | Regulates transcription through chromatin remodeling | Not known |
| CIC | Q96RK0 | 50 | 22 | 0.27 | 10.27 | 1.09 | S277 | 0.62 | 9.8 | >12 | Linear | MAPK | MAPK | PMID: 33103082 | ++ | Member of the high mobility group (HMG)-box superfamily of transcriptional repressors. Represses DUSP6 expression | Induces binding of 14-3-3 proteins, which blocks its normal inhibition of transcription. |
| ATXN1L | P0C7T5 | 17 | 5 | 0.22 | 10.72 | | S284 | 0.56 | 6.0 | 6.1 | Linear | RSK | | PMID: 29578365; PMID: 28178529 | +++ | Part of the ATXN1L-CIC transcriptional repression complex that binds chromatin and blocks transcription | Induces binding of 14-3-3 proteins |
| TAF3 | Q5VWG9 | 15 | 2 | 0.14 | 8.60 | | S199 | 0.43 | 10.0 | 6.5 | Linear | RSK | | PMID: 33795473 | + | Part of the TFIID transcription factor protein complex that plays a central role in mediating promoter responses to variou | Not known |
| AKAP1 | Q92667 | 12 | 2 | 0.10 | 10.79 | 0.86 | S151 | 0.84 | 6.0 | >12 | Bi-phasic | MAPK | PKC | PMID: 16669629 | - | Binds to type I and type II regulatory subunits of PKA and anchors them to the mitochondrion. | Induces dissociation of PP1 phosphatase |
| TSC2 | P49815 | 24 | 6 | 0.12 | 11.17 | 1.04 | S1132 | 0.95 | 3.2 | 6.1 | Linear | RSK | AKT | PMID: 12172553 | + | Suppresses mTOR signaling | Induces dissociation from TSC1, activating S6 Kinase 1 (RPS6KB1) |
| CDC25C | P30307 | 6 | 2 | 0.11 | 11.44 | 1.15 | T48 | 0.65 | 4.5 | 5.4 | Linear | MAPK | CDK1 | PMID: 7513216; PMID: 20668692 | - | Phosphoprotein phosphatase involved in regulating cell cycle progression | Activates its protein phosphatase activity, allowing cell cycle progression |
| ABL1 | P00519 | 16 | 3 | 0.13 | 8.71 | 1.21 | S718 | 0.54 | 2.6 | >12 | Bi-phasic | RSK | JNK | PMID: 15696159 | +++ | Coordinates actin remodeling through tyrosine phosphorylation of proteins controlling cytoskeleton dynamics. | 14-3-3 binding site. Causes sequestration to the cytoplasm |
| PLEKHG3 | A1L390 | 34 | 11 | 0.23 | 9.36 | 1.19 | S962 | 0.31 | 6.3 | 7.3 | Bi-phasic | MAPK | | PMID: 27555588 | - | Rac/Rho GEF. Involved in regulation of cell migration and establishment of cell polarity. | Not known |
| MAST4 | O15021 | 50 | 6 | 0.17 | 7.95 | | S1779, S2552 | 0.34/0.17 | 2.8-3.6 | 7-12 | Linear | MAPK/RSK | | PMID: 35803931 | +++ | A member of the microtubule-associated serine/threonine protein kinase family. Function unclear, but could regulate st | Not known |
| ARHGAP21 | Q5T5U3 | 52 | 10 | 0.24 | 11.13 | 1.80 | S1713 | 0.40 | 8.2 | 6.7 | Linear | RSK | | PMID: 29856495 | - | Functions as a GTPase-activating protein (GAP) for RHOA and CDC42. Thought to be involved in cytoskeletal remodeling. | Not known |
| SHROOM1 | Q2M3G4 | 15 | 4 | 0.14 | 9.21 | | S188 | 0.46 | 2.6 | >12 | Linear | RSK | | PMID: 27758857 | ++ | Control cell shape by regulating cytoskeletal dynamics through binding F-actin and ROCK | Induces binding of 14-3-3 proteins at the ASD1 F-Actin binding domain |
| SHROOM3 | Q8TF72 | 44 | 15 | 0.20 | 9.70 | 2.13 | S816 | 0.29 | 3.8 | 5.9 | Linear | RSK | | PMID: 27758857 | - | Control cell shape by regulating cytoskeletal dynamics through binding F-actin and ROCK | Not known |
